# LiMMBo: a simple, scalable approach for linear mixed models in high-dimensional genetic association studies

**DOI:** 10.1101/255497

**Authors:** Hannah Verena Meyer, Francesco Paolo Casale, Oliver Stegle, Ewan Birney

## Abstract

Genome-wide association studies have helped to shed light on the genetic architecture of complex traits and diseases. Deep phenotyping of population cohorts is increasingly applied, where multi-to high-dimensional phenotypes are recorded in the individuals. Whilst these rich datasets provide important opportunities to analyse complex trait structures and pleiotropic effects at a genome-wide scale, existing statistical methods for joint genetic analyses are hampered by computational limitations posed by high-dimensional phenotypes. Consequently, such multivariate analyses are currently limited to a moderate number of traits. Here, we introduce a method that combines linear mixed models with bootstrapping (LiMMBo) to enable computationally efficient joint genetic analysis of high-dimensional phenotypes. Our method builds on linear mixed models, thereby providing robust control for population structure and other confounding factors, and the model scales to larger datasets with up to hundreds of phenotypes. We first validate LiMMBo using simulations, demonstrating consistent covariance estimates at greatly reduced computational cost compared to existing methods. We also find LiMMBo yields consistent power advantages compared to univariate modelling strategies, where the advantages of multivariate mapping increases substantially with the phenotype dimensionality. Finally, we applied LiMMBo to 41 yeast growth traits to map their genetic determinants, finding previously known and novel pleiotropic relationships in this high-dimensional phenotype space. LiMMBo is accessible as open source software (https://github.com/HannahVMeyer/limmbo).

**Author summary:** In multi-trait genetic association studies one is interested in detecting genetic variants that are associated with one or multiple traits. Genetic variants that influence two or more traits are referred to as pleiotropic. Multivariate linear mixed models have been successfully applied to detect pleiotropic effects, by jointly modelling association signals across traits. However, these models are currently limited to a moderate number of phenotypes as the number of model parameters grows steeply with the number of phenotypes, raising a computational burden. We developed LiMMBo, a new approach for the joint analysis of high-dimensional phenotypes. Our method reduces the number of effective model parameters by introducing an intermediate subsampling step. We validate this strategy using simulations, where we apply LiMMBo for the genetic analysis of hundreds of phenotypes, detecting pleiotropic effects for a wide range of simulated genetic architectures. Finally, to illustrate LiMMBo in practice, we apply the model to a study of growth traits in yeast, where we identify pleiotropic effects for traits with formerly known genetic effects as well as revealing previously unconnected traits.

## Introduction

Quantitative measurements of organisms have been a bedrock of genetics since the birth of this science [1]. One can measure many different traits of an organism and it is natural to want to jointly analyse the measurements to discover new genetic associations, generically called multi-trait analysis. Many cohort studies today have rich, high-dimensional datasets ranging from studies in model organisms such as yeast and *Arabidopsis thaliana* to human molecular, morphological or imaging derived traits [2–6]. Genetic association studies of high-dimensional traits have often used approximate methods such as dimensionality reduction of the phenotypes prior to the genetic mapping and univariate trait-by-trait analyses followed by post-hoc integration using meta-analysis. Although both strategies have been successful in a variety of scenarios [7–10], neither uses the potential of genetics to inform the joint multi-trait analysis. In contrast, multivariate regression models explicitly model the genotypic association across multiple phenotypes.

A recent innovation in regression models for genetics has been to use linear mixed models (LMMs). LMMs in genetics use the random effect term for explicitly modelling complex genetic structure [11, 12] thereby retaining calibrated test statistics even in structured populations. LMMs are common in genetic studies of animal and plant breeding, and more recently have been applied to account for population structure and relatedness in human populations [10, 13, 14]. In addition, they allow for estimating narrow sense heritability [15, 16].

Most recently, extensions of the classical LMM have enabled joint genetic analyses of multiple traits, where the random effect component captures both the genetic covariance between individuals (sample-to-sample) and the traits (trait-to-trait) [17–19], accounting for the genetic trait covariance. Additionally, these models also consider a residual trait covariance, which capture covariances between traits not due to genetics. Importantly, while the genetic sample covariance can be directly estimated from the genetic variation data itself, e.g. using a realised relationship or identity by descent, the trait covariance matrices need to be statistically estimated, typically using a (restricted) maximum likelihood procedure.

Current multi-trait LMM implementations scale reasonably well with the number of analysed samples [18, 19] but not as well with number of the traits.Specifically, computations become prohibitive as soon as a few tens of traits (P) are considered, with a computational complexities ranging O(P5) to up to O(P7) for existing methods [17, 19].

Here we developed a simple, but surprisingly effective heuristic to efficiently estimate large trait covariance matrices in linear mixed models using bootstrapping (LiMMBo), thereby allowing for the analysis of datasets with a large number of phenotypic traits. We first validate LiMMBo through comparisons to exact inference methods on a smaller and tractable datasets, demonstrating the consistency of the variance component estimates obtained using the model.

Importantly, LiMMBo is faster than existing methods, even for moderately sized problems. This enables both efficient variance component estimation but also genetic mapping for high-dimensional phenotypes with up to hundreds of traits. Using simulations, we show that the ability to scale multi-trait LMMs to large numbers of traits substantially increases the power to detect genetic effects. Finally, we illustrate the effectiveness of LiMMBo through application to a yeast quantitative trait loci study [2] with 41 measurements. We observed an increase in power compared to univariate models and used LiMMBo to explore the pleiotropic relationship between traits.

## Results

### Covariance estimation via bootstrapping

LiMMBo builds on a multivariate LMM framework with multiple fixed and random effect components [19, 20]. This model can be used to estimate genetic and non-genetic variance components, and it has been applied to test for genetic associations between individual variants and moderately sized groups of traits [21, 22]. Briefly, this LMM model can be cast as

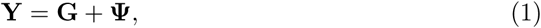

where the *N* | *P* phenotype matrix **Y** for *N* individuals and *P* traits is modelled as the sum of a genetic (or polygenic) component **G** and a noise component **Ψ**. We have omitted additional fixed effects for notational brevity. Here, **G** and **Ψ** are random effects following matrix-normal distributions:

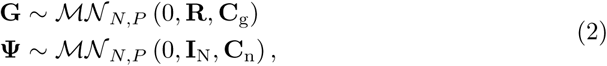

where **R** denotes the *N* × *N* genetic relationship matrix, **I**_N_ is the *N* × *N* identity matrix and **C**_g_ and **C**_n_ are the genetic and the residual trait *P* × *P* covariance matrices, respectively. The marginal likelihood of the model in equation Eq. 2 can be expressed in terms of a multivariate normal distribution of the form

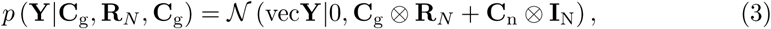

where the covariance structure of the phenotypes (in shape of the *N* × *P* phenotype vector vec (**Y**) through stacking the columns of the phenotype matrix) is described by the sum of the Kronecker products ⊗ of the sample and trait covariance terms. This model enables efficient inference schemes by exploiting Kronecker identities for the eigendecomposition of the full covariance matrix [18–20, 23]. In particular, it allows for decoupling the decomposition of **C**_g_ and **R***_N_*, which greatly increase the efficiency of the inference as **R**_N_ is constant. The model in Eq. (3) also corresponds to the null model when using the multi-trait LMM for genetic association mapping. While there exist exact schemes for refitting the variance components for each variant tested [18], approximations that keep the fitted variance components **C**_g_ and **C**_n_ from the null model constant across tests frequently yield satisfactory results [12, 17, 20].

The complexity of a multivariate LMM implementation previously described in [19, 20] (herein ‘standard REML’) is *O*(*N*^2^ + *t*(*N P*^4^ + *P*^5^)) with *N* samples, *P* traits, and *t* iterations of Broyden’s method, which uses an approximation of the second derivative for optimising the REML of the parameter estimates. From this scaling behaviour, it becomes evident that as the number of traits increases, the complexity increases steeply (as is the case for other inference schemes, Supplementary Tab. S1), thereby limiting applications for larger numbers of traits.

The key innovation in LiMMBo is to perform the variance decomposition on *b* bootstrap samples of a subset of *s* traits, and then to appropriately use those bootstrap samples to reconstruct the full **C**_g_ and **C**_n_ matrices (Fig. 1, right panel). By breaking down the computationally expensive variance estimation step into *b* independent steps, LiMMBo reduces the overall complexity of the ‘standard REML’ to *O*(*N*^2^ + *bt*_1_(*N s*^4^ + *s*^5^) + *t*_2_*P*^2^). Parameter estimation for the full trait set is only required for the final covariance matrices **C**_g_ and **C**_n_ and is achieved by finding the best fit of the bootstrap estimates to a *P* × *P* covariance matrix (see Methods). While this approach keeps the complexity at *O*(*P*^2^), it has the additional advantage of allowing for trivial parallelisation of the covariance estimation step. The variance decomposition of each bootstrap is computed independently and our implementation allows for making use of multiple cores.

**Fig 1.**
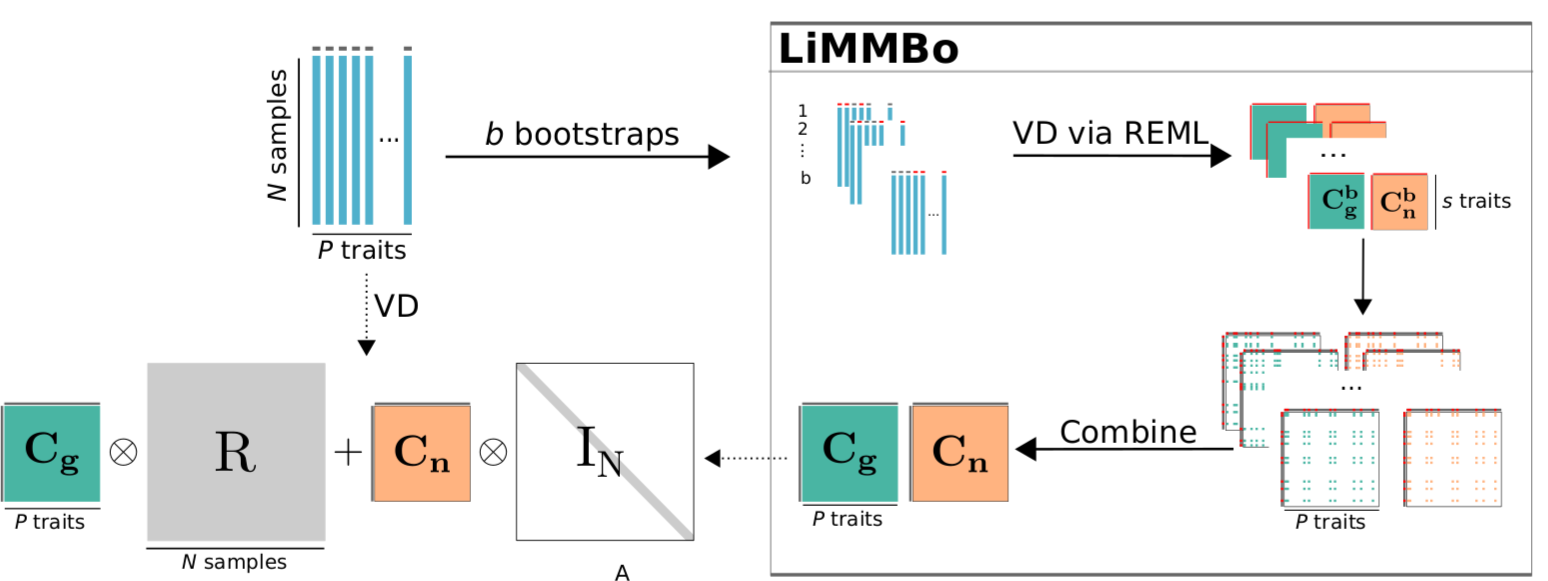
Variance decomposition with REML and LiMMBo. REML approach (left): the phenotype set of *P* traits and *N* samples is decomposed into its *P* × *P* trait-to-trait covariances **C**_g_ and **C**_n_, based on the provided *N* × *N* genetic sample-to-sample kinship estimate matrix **R**. The noise sample-to-sample matrix **I** is assumed to be constant (identity matrix). Existing implementations of LMM to fit such variance decomposition (VD) models via REML are limited to moderate numbers of phenotypes. LiMMBo approach (right): for higher trait-set sizes, the phenotypes’variance components are estimated on *b s*-sized subsets of *P* which are subsequently combined into the overall *P* × *P* covariance matrices **C**_g_ and **C**_n_.

### Scalability

To assess the scaling of LiMMBo, we fit the model to simulated datasets with increasing numbers of phenotypes. Fig. 2 shows both the overall compute time and a break down into the two main components of the method, bootstrapping and the combination of the bootstrapping results. The majority of compute time is needed for the variance decomposition of the bootstrapped subsets, which can be trivially parallelised across bootstraps. As a comparison, the time taken by the standard REML approach quickly exceeds the time of LiMMBo and becomes infeasible for more than 30 traits.

### LiMMBo yields reliable covariance estimates

To assess the accuracy of LiMMBo for covariance estimation of the trait covariances **C**_g_ and **C**_n_, we considered in-silico simulations with known ground truth. We simulated datasets with different extents of population structure (based on genotype data of the 1000Genomes Project [24], see Methods), varying the extent of the genetic population effect and different numbers of traits.

First, we compared the LiMMBo and the standard REML fits in terms of their accuracy to recover the true simulated covariance matrix. This comparison is only feasible in the regime of low to moderate number of traits (i.e., lower than 30) where existing REML implementations can be applied. Reassuringly, we observed that both approaches provide consistent estimates of comparable accuracy, independent of the number of traits (Fig. 3A).

**Fig 2.**
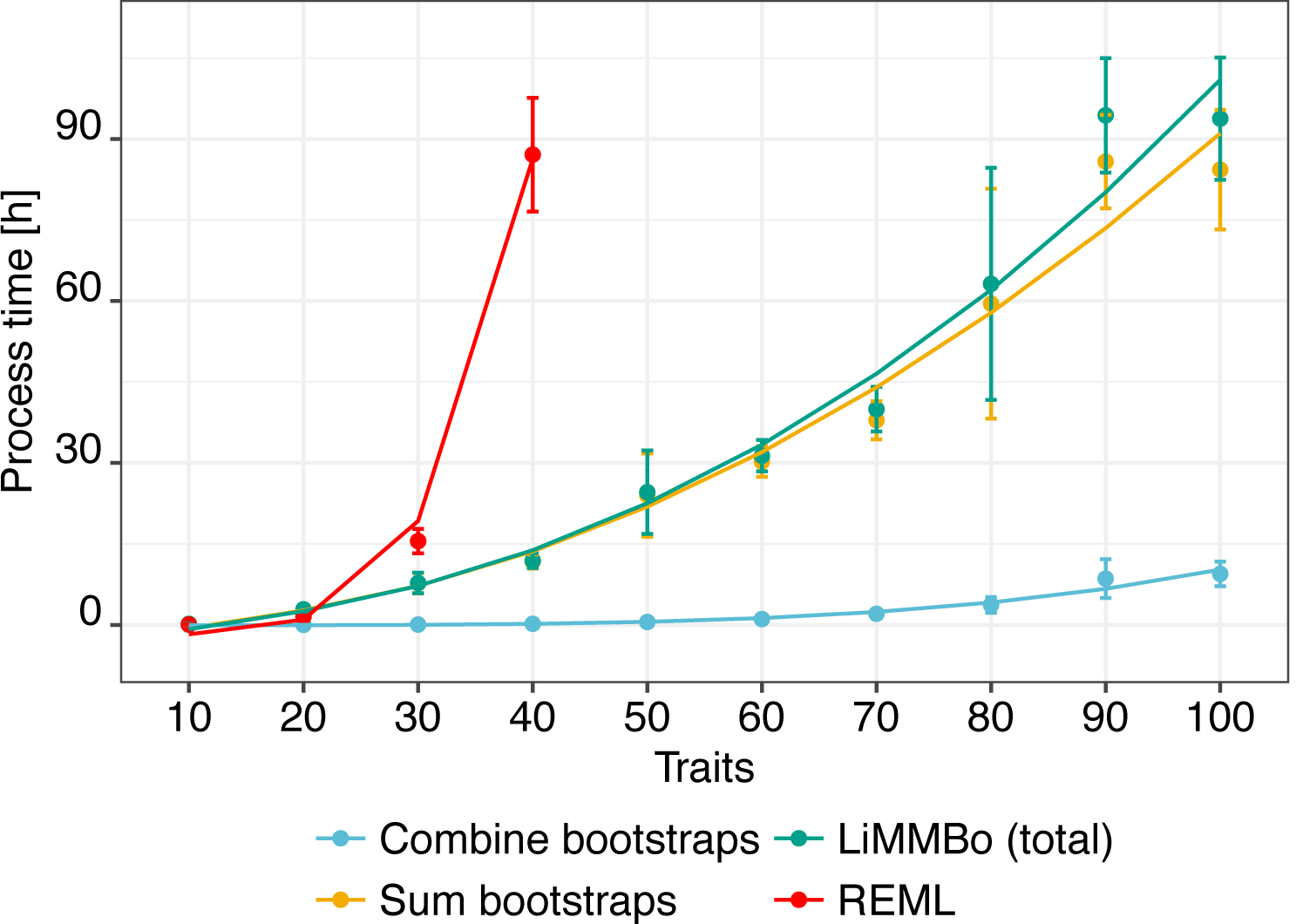
Scalability of REML and LiMMBo. Empirical run times for standard REML inference and LiMMBo on ten simulated datasets with *N* = 1,000 individuals, increasing number of traits (*P* ∈ {10, …, 100} and different amount of variance explained by the genetics (*h*_2_ ∈ {0.2,0.5,0.8}). Shown is the average empirical run time for 30 experiments per trait size (10 per *h*_2_), with error bars denoting plus or minus one standard deviation across experiments. Lines denote a fit of the theoretical complexity to the observed run times: bootstrapping step (orange): *b*(*N s*^4^ + *s*^5^); the combination of the bootstrapping (blue): *P*^2^, their combined run time (turquoise): *b*(*N s*^4^ + *s*^5^) + *P*^2^ and the standard REML approach (red): *N P*^4^ + *P*^5^. *b*: number of bootstraps, *s*: bootstrap size, *P*: phenotype size, *N*: sample size. The majority of the run time is allocated to the bootstrapping. Run times for the standard REML are depicted up to *P* = 40 when they already exceed the run times for *P* = 100 in the LiMMBo approach.

Next, we assessed the utility of LiMMBo estimates of the trait covariance matrices for carrying out multi-trait GWAS. Specifically, we assessed the calibration of type-I error rates using phenotypes simulated from the null (no causal variant) and applying LiMMBo to fit the the trait covariances **C**_g_ and **C**_n_ which we then used in a multivariate LMM [17]. For low to moderate number of traits we compared the calibration to a multi-trait LMM using standard REML-derived estimates of **C**_g_ and **C**_n_. Association tests based on the random effect covariance estimates using both inference schemes were calibrated across varying proportion of variance explained by genetic effects and different numbers of traits, including the regime of large *P*, where existing methods cannot be applied (Fig. 3B).

In principle, multivariate genetic analyses in higher dimensions are also possible using simple linear models without a random effect component, thereby avoiding the computational bottleneck of genetic trait covariance estimation. However while population structure can be accounted for within this approach, for example by including principal components of the genotypes as covariates into the model, these methods have known limitations when the individuals are related [12, 19]. Consistent with this, we found that the linear model was poorly calibrated in such structured populations (Supplementary Tab. S4), clearly demonstrating the strength of the mixed-model approach LiMMBo.

### Multi-trait genotype to phenotype mapping increases power for high-dimensional phenotypes

First, we consider simulated data to assess whether the power benefits of multivariate LMMs translate to high-dimensional phenotypes when using a fixed effect test with as many degrees of freedoms as traits. We examined a wide range of simulation settings, simulating up to 100 traits and varying proportions of traits affected by genetic variants. The mean genetic variance across all traits was kept constant (i.e with an increase in the affected traits the contribution of the genetic component per trait decreases). For each set-up, we simulated 50 different phenotypes and estimated the trait covariance matrices **C**_g_ and **C**_n_ via LiMMBo.

We used these estimates in a multivariate LMM to test the association between the known causal SNPs (from simulation) and the phenotypes and compared them to results of univariate LMMs of the causal SNPs and the phenotypes. Fig. 4 compares the power of the multivariate and univariate LMM as the percentage of significant SNPs out of all true causal SNPs (univariate p-values were adjusted for multiple testing across traits, see Methods). In the scenario where all traits are affected by the fixed genetic effects, the burden of multiple testing in the univariate models weighs at least as heavy as the increased number of degrees of freedom. For the highest number of traits simulated both models are comparable in the number of causal SNPs they detect. For the other trait sizes tested, the multivariate model out-performs the univariate model by far (Fig. 4A). The advantage of the multivariate model to exploit correlated trait structures becomes evident when the proportions of traits affected by the causal SNPs is increased. The univariate model suffers from the weaker genetic components when a large number of traits is affected and loses power. In contrast, the multivariate model can still detect increasing percentages of true causal SNPs (Fig. 4B). For both the univariate and multivariate model, the number of detected SNPs decreases with increasing variance explained by genetics, as the effect sizes of the SNPs (fixed for all simulations) become negligible compared to the overall genetic variance. However, the multivariate model is still able to exploit the correlation of the SNP effects across traits and detects more SNPs in cases of high genetic background structure (Fig. 4C). An overview of all parameter comparisons can be found in Supplementary Fig. S2.

**Fig 3.**
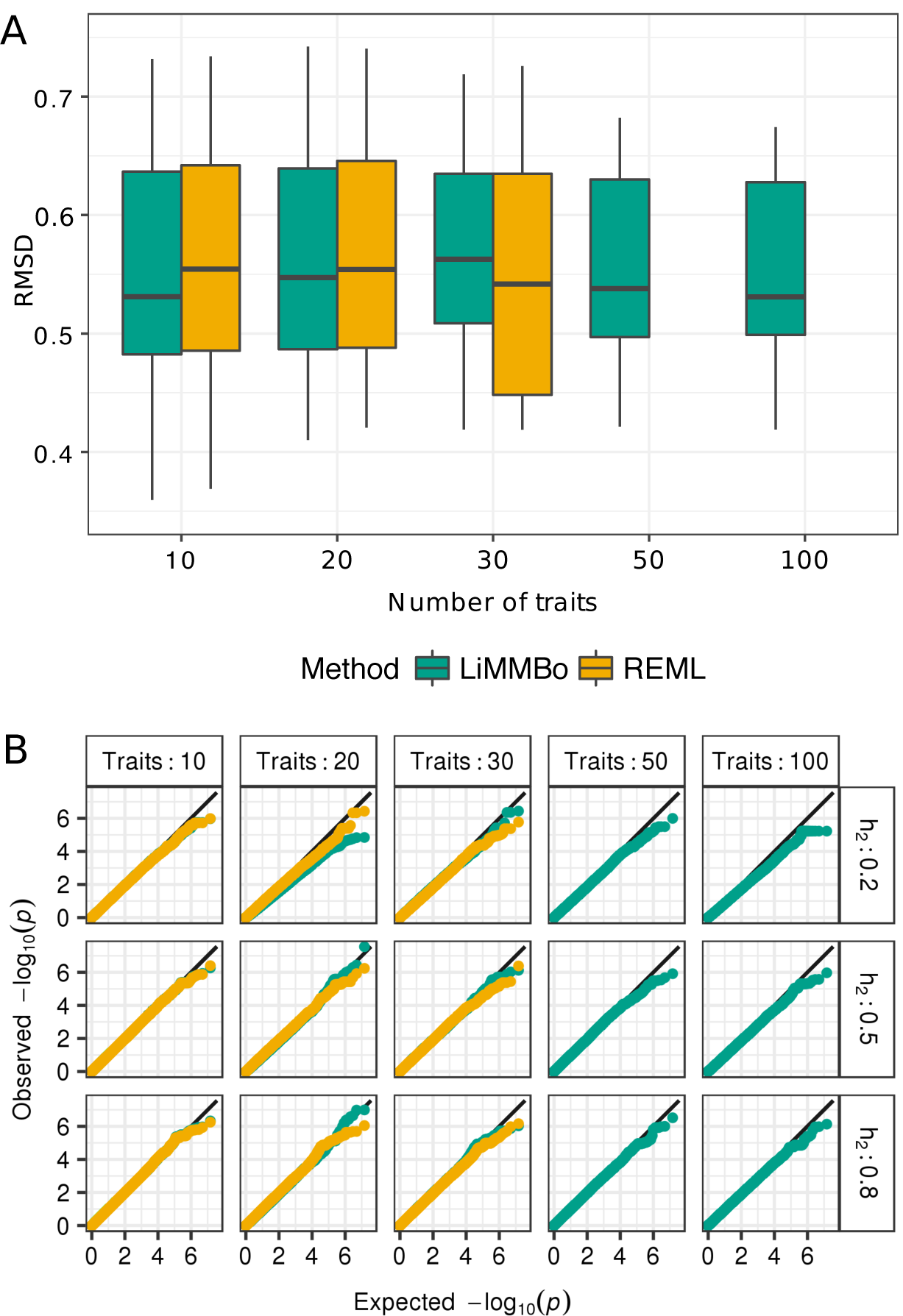
Comparison of trait-by-trait covariance estimates derived using standard REML and LiMMBo. Phenotypes with increasing percentages of variance explained by genetic effects (*h*_2_ ∈ {0.2,0.5,0.8}) and for increasing numbers of traits were simulated. The genetic and noise trait covariance matrices **C**_g_ and **C**_n_ were estimated using both LiMMBo and standard REML. **A** Comparison of the estimated trait covariances to the simulated (’true’) matrices using the root mean squared deviation (RMSD). For each of the simulation scenarios, ten independent datasets generated for each setting. Boxplots show the distribution of RMSD values for **C**_g_ and **C**_n_ across the simulation settings. For moderate trait set sizes ranging from 10 to 30 traits, LiMMBo and conventional REML yield consistent covariance estimates. RMSD in covariance estimates via LiMMBo remained stable for larger numbers of traits (*P* ∈ {50,100}). **B** Comparison of model calibration in multi-trait GWAS with LiMMBo and standard REML derived covariance matrices. Statistical calibration of p-values under the null model was assessed for each of the trait sizes and percentages of variance explained by genetic effects. Quantile-quantile plots show uniform distribution for both methods across all trait sizes and levels of proportion of variance explained by genetics. Multi-trait GWAS for phenotypes with 50 and 100 traits was only possible with LiMMBo.

**Fig 4.**
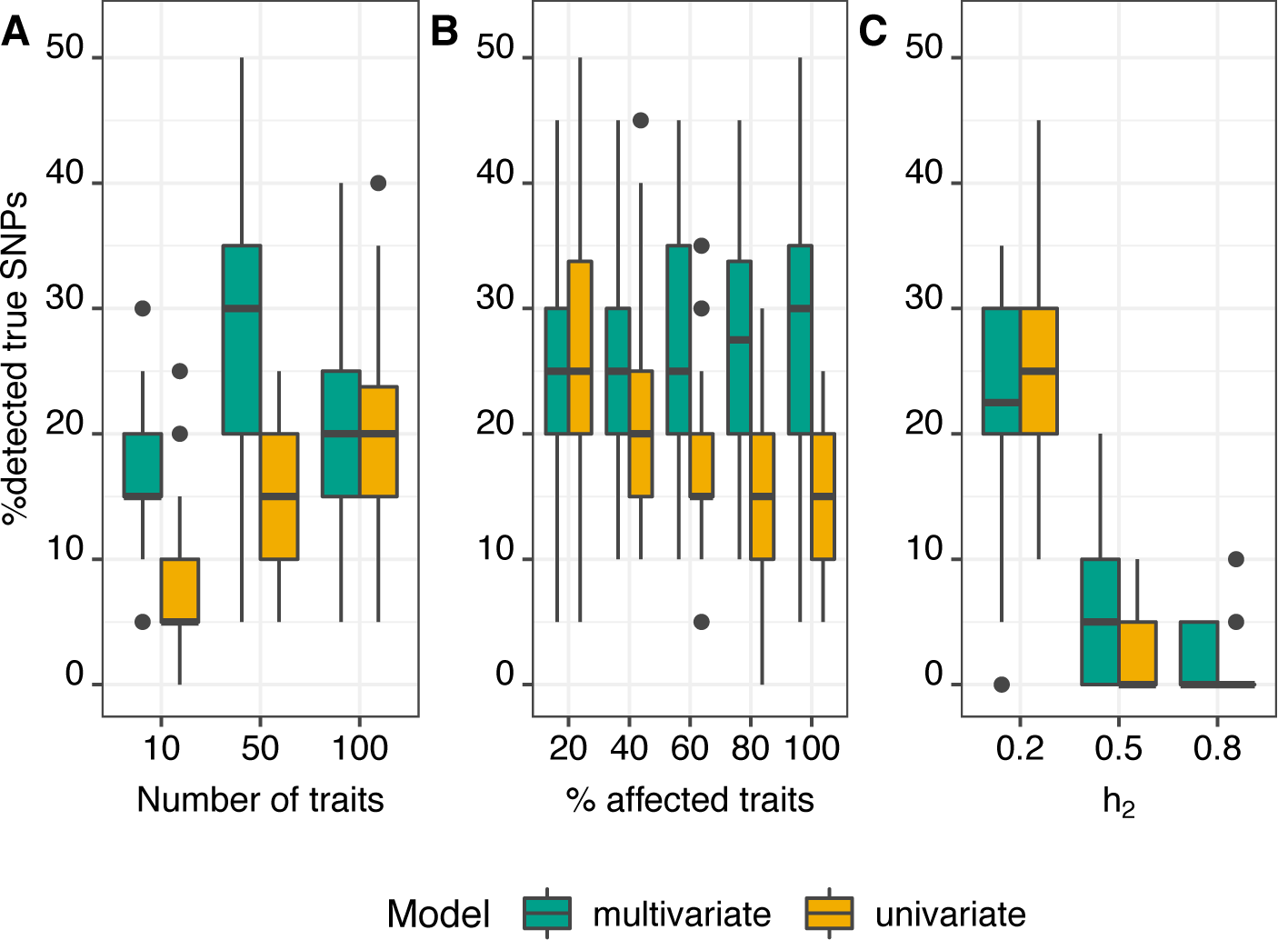
Power comparison for multivariate and univariate LMMs of high-dimensional phenotypes. Each panel shows the influence of one simulation parameter on the power to detect the causal SNPs. When investigating one parameter, the other parameters were fixed at a certain value. For each set-up, 50 independent datasets were simulated and analysed. A. Influence of the number of traits: proportion of traits affected and the total genetic variance fixed at *a* = 1 and *h*_2_ = 0.2, respectively. B. Influence of proportion of traits affected: trait size and total genetic variance fixed to *P* = 50 and *h*_2_ = 0.2 respectively. C. Influence of total genetic variance: trait size and proportion of traits affected fixed to *P* = 100 and *a* = 0.6.

### Multi-trait GWAS in yeast

Next, to illustrate the utility of LiMMBo, we applied the model to a QTL dataset of growth rates measured in a yeast cross cross for 41 different conditions [2], which is a well-placed complexity of traits given our simulations. Fig. 5A depicts results from a multi-trait GWAS based on LiMMBo covariance estimates, as well as a single-trait GWAS analysis, where each trait was analysed separately. Multi-trait GWAS with LiMMBo (blue) detected more significant associations, identifying 384 associations versus 275 associations (accounting for linkage disequlibriium, *r*^2^ < 0.8 in a 3kb window; see below) in the single-trait GWAS (orange; minimum p-value per SNP across all 41 single-trait GWAS, adjusted for multiple testing). This increase in detected associations was consistent across different ranges of LD criteria to define individual loci (Supplementary Tab. S5), with on average 29% more significant loci in the multi-trait GWAS. In contrast there are a few significant loci for the single-trait GWAS where the multi-trait analyses either does not reach significance (e.g. on chromosome 7) or does not detect any association (e.g. on chromosome 4). For these loci, the underlying genetics seem to be trait specific to magnesium sulfate and hydroquinone, respectively (Supplementary Fig. S5.

In addition to the significance of an association, linear mixed models provide effect size estimates for the tested SNPs. By analysing the effect sizes of significantly associated SNPs across traits, we can explore pleiotropic effects of these significant loci. We limited the analysis to the multi-variate effect size estimates from significant variants located within a gene body to ensure we could link the variants to specific genes for downstream analysis. Subsequently, we pruned these variants for LD (*r*^2^ > 0.2, 3kb window; see methods) to remove potential bias in the clustering due to an over-representation of variants from large loci.

Most of these resulting 210 loci have strong effect size estimates for more than one trait, i.e., most loci seem to be pleiotropic (Fig. 5B, non-zero effect sizes in columns). Furthermore some traits have striking contributions from multiple loci across the genome, in particular xenobiotic growth conditions e.g. zeocin [25] and neomycin [26] (Fig. 5B, large effect sizes across rows).

To gain more insights into the trait architecture, we analysed the effect size estimates across loci and traits using hierarchical clustering (robustness of the clustering assessed via a bootstrapping-based method (pvclust, [27])). Overall, we observed 14 stable trait and 136 stable variant clusters, many of which are nested; these clusters collapsed into 6 trait and 19 variant exclusive groupings at the highest branch (Supplementary Fig. S3).

These six significant clusters for traits (Fig. 5B, blue row dendrogram) include classically linked carbon metabolism sources (lactose, lactate and ethanol), and other clusters which there is literature support for. For example, there is consistent gene expression changes upon treatment such as DNA replication agents hydroxyurea and 4-nitroquinoline-l-oxide (x4NQO) [28] or trehalose and sorbitol which have previously been shown to have synergistic effects on viability in yeast [29]. For other clusters, such as SDS and Hydroxybenzaldehyde or magnesium sulfate and berbamine we were unable to find literature support but this could be a candidate clustering of these growth phenotypes for further investigation. Out of the 19 stable SNP clusters (Fig. 5B, blue column dendrogram), many clusters include disjoint regions across a chromosome and twelve clusters (Fig. 5B, grey boxes) span multiple chromosomes. Some of the clusters extending across chromosomes have suggestive common annotation, such as *cluster a* which has two members of the nuclear pore complex (NUP1, NUP188), and *cluster b* with a common set of vesicle associated genes (ATG5, PXA1,VPS41; Fig. 5B, labelled boxes). The small size of clusters inhibited any systematic gene ontology based enrichment, but the ability to explore both multiple traits and multiple loci from the multi-trait GWAS provides stimulating hypothesis generation.

**Fig 5.**
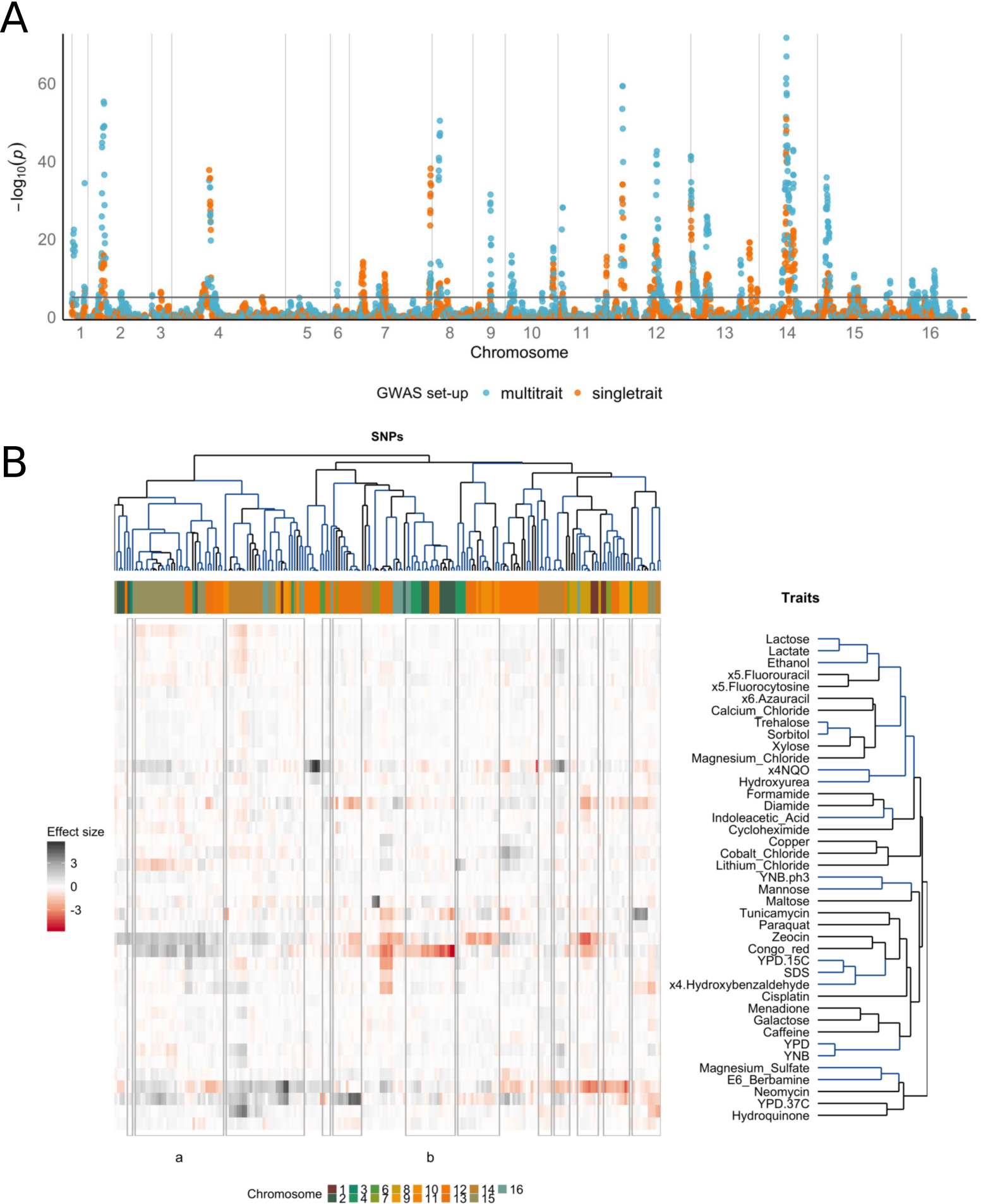
GWAS across 41 yeast growth phenotypes. A. Manhattan plot of p-values from single-trait and multi-trait GWAS. The single-trait GWAS p-values were adjusted for multiple testing and only the minimum adjusted p-values across all 41 traits per SNP are shown. The significance line is drawn at the empirical FDR_stGWAS =_ 8.6 × 10*^−6^* **B. multi-trait GWAS effects size estimates.** Effect size estimates of LD-pruned (3kb window, *r*^2^ > 0.2), significant SNPs located within a gene were clustered by loci and traits (both hierarchical, average-linkage clustering of Pearson correlation coefficients). Traits and SNPs in stable clusters (pvclust *p* < 0.05) are marked in blue. Grey boxes indicate stable SNP clusters spread across at least two chromosomes; a and b label two clusters for which suggestive common annotation was found, for details see text.

Finally, for comparison, we repeated the same clustering analysis based on the results from the single-trait model. This identified only 9 stable traits and 117 clusters (Supplementary Fig. S4,S6), compared to 14 and 136 clusters from the multi-trait analysis. As the number of input significant variants differ substantially between the two analyses, it is complex to directly compare the two clusterings. However, both the higher number of variants and the larger number of clusters shows that there is a clear advantage in the multi-trait analysis.

## Discussion

We have developed LiMMBo, a new methods for estimating trait-covariances for multi-trait LMMs, which offers substantially enhanced scalability using a bootstrap strategy. The most important benefit of LiMMBo is the scalability to 100s of phenotypes, both because of its sub-sampling method and due the practical aspect that major aspects of the computation can be parallelised. Our implementation detects multi-threaded cores automatically, thereby taking advantage of this opportunity. In practice, this means that trait sizes up to 30 or 40 can be run in hours, rather than taking several days when using existing, full maximum likelihood methods for inference. Complex datasets of up to 100 traits, which cannot be analysed using existing implementations, are tractable using LiMMBo. In simulations, we show that the resulting covariance matrices are as good an estimator of the real covariance matrices as the maximum likelihood methods, yielding well-calibrated test statistics when used for genetic association analyses in LMMs. We applied LiMMBo to a multi-trait yeast dataset [2], showing an increase of power for loci discovery, in particular pleiotropic loci, and the ability to analyse the pleiotropic effects of each locus. LiMMBo is accessible as an open source, python module (https://github.com/HannahVMeyer/LiMMBo) compatible with the LIMIX package [20, 30] for LMMs.

Much of the attraction of linear mixed models in genetics has been their ability to model complex genetic relatedness. Unsurprisingly, simple linear models are not suitable for analysing phenotypes with complex underlying genetic relatedness, whereas LMMs with the covariance matrices estimated by LiMMBo are appropriate and possible up to 100s of traits. Complex relatedness in populations is widespread in plant‐ and animal breeding in particular [10, 13], and increasingly common in human bottleneck populations [14]. Furthermore, as the population numbers increase in human genetics, complex cryptic relationship structures are more prevalent [31], meaning that methods such as LiMMBo will be more applicable in the future in human genetics.

The robust estimation of large trait covariance matrices is a recurrent statistical challenge in genomics, from statistical genetics to single-cell analysis. The ability to accurately estimate large trait covariance matrices using this bootstrap method may be applicable to domains other than GWAS, e.g. many gene expression studies use covariance matrices. Previous work from Schaefer and colleagues [32] showed the large gene dimensions coupled with small(er) sample sets meant that empirical covariance matrices could not be accurately estimated; other investigators [33–35] used shrinkage methods to create valid covariance matrices. The work from Teng and colleagues [36] uses subsampling but with strong shrinkage priors to generate the final covariance matrix. By fitting the bootstrap average to the closest true covariance, LiMMBo ensures positive-semidefiniteness of the covariance while avoiding ill-conditioned matrices, which usually introduces large biases in the final use of these models. We are actively exploring the use of the LiMMBo covariance estimation in this and other areas.

Our ability to generate large cohorts of well phenotyped and genotyped individuals has forced the development of many new methods in statistical genetics. With the advent of genotyped human cohorts up to 500,000 individuals with over 2,000 different traits [37], and plant phenotyping routinely in the 1,000s of individuals from structured crosses with 100s to 1,000s of image based phenotypes [3, 13], we need both informative and scalable methods. LiMMBo extends the reach of linear mixed models into this new regime, allowing for new complex genetic associations to be made and a more informative exploration of the underlying biological effects.

## Materials and Methods

The LiMMBo implementation and all analyses scripts can be found at https://github.com/HannahVMeyer/limmbo and https://github.com/HannahVMeyer/LiMMBoAnalysis

### LiMMBo implementation

LiMMBo is implemented as an open-source python module based on the *limix* (version:limix-v1.0.12, url:https://github.com/limix/limix) module of the LIMIX framework for linear mixed models [20].

#### Covariance estimation

Covariance estimation via LiMMBo can conceptually be separated into three steps: i) division of the full dataset into subsets, ii) variance decomposition on the subsets using the REML approach implemented in [19, 20] and iii) combine the results obtained from the subsets.

In detail, from the total phenotype set with *P* traits, *b* subset of *s* traits are randomly selected. *b* depends on the overall trait size *P* and the sampling size *s* and is chosen such that each two traits are drawn together at least *c* times (default: 3).

For each subset, the variance decomposition is estimated via *mtSet.pycore.modules.multiTraitSetTest* and *.fitNull* i.e. the null model of the mvLMM (Eq. 1). For each bootstrap, the *s* × *s* covariance matrices 
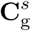
 and 
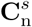
 are recorded.

The challenge lies in combining the bootstrap results in a way, that the resulting **C**_g_ and **C**_n_ matrices are covariance matrices i.e. positive semi-definite, and serve as good estimators of the true covariance matrices. First, the covariance estimates for each trait pair are averaged over the number of times they were drawn. The covariance estimates of the *b* subsets are then combined by a least-squares fit to the closest positive-semidefinite matrices via the gradient-based low-memory Broyden-Fletcher-Goldfarb-Shanno algorithm (BFGS) [38, 39] implemented in *fmin l bfgs b* of the SciPy python library [40]. The average covariance estimates are used to initiate the optimisation matrices via *limix.CFreeFormCF* and *.setParamsCovariance*. The model makes use of a Cholesky decomposition of the initial matrices to be fitted via *.getParams()*, resulting in 1/2 *P* (*P* + 1) model parameters to be fitted. **C**_g_ and **C**_n_ are fitted separately.

#### Genetic association testing with LiMMBo

Genetic association testing with LiMMBo is split into two parts. First, the full trait covariance matrices of the genetic **C**_g_ and non-genetic **C**_n_ random effects are estimated via the covariance estimation scheme outlined above. Second, the covariance estimates are used as input parameters for the multivariate linear mixed model with the full phenotype set as the response variable, the genetic variant of interest as fixed effect and possible, further non-genetic covariate effects (Eq. 5).

### Data simulation

#### Genotypes

Genotypes were simulated similar to strategies described in [19, 41]. A cohort of 1,000 synthetic genotypes were generated based on real genotype data from four European ancestry populations of the 1,000 Genomes (1KG) Project (populations: CEU, FIN, GBR, TSI) [24]. Each newly synthesised individual is assigned to a specified number of ancestors *a* from the original 1KG Project and their genome split into blocks of 1,000 SNPs (thereby retaining realistic LD structure between SNPs). For each SNP block, the ancestor is randomly chosen and its genotype is copied to the individuals genome. Choosing a low number of ancestors for the simulation introduces relatedness among individuals, while high numbers lead to unstructured populations. Two cohorts were simulated, one with a low number of ancestors (*a* = 2, *popStructure*) and one with a high number of ancestors (*a* = 10, *noPopStructure*). The latter is only used for the comparison of calibration of the multivariate linear model to the multivariate linear mixed model. The resulting population structures are depicted in Supplementary Fig. S1.

#### Phenotypes

Phenotypes were generated as the sum of up to four components: i) genetic variant effects, ii) infinitesimal genetic effects (based on population structure), iii) covariate effects and iv) observational noise via [42]. All components have a percentage of common effect, i.e. the effect was shared across traits and a specific effect. A list and description of all simulation parameters can be found in Supplementary Tab. S2. For each of the genotype cohorts described above, different phenotype scenarios depending on percentage of variance explained by genetics *h*_2_ and number of traits *P* were simulated: i) *h*_2_ = {0.2,0.5,0.8} and ii) *P* = {10,20, …, 100}. The variance explained by every components was fixed by scaling each of the components with a factor *a* such that their average column variance 
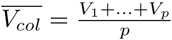
 explains a fraction *x* of the total variance: 
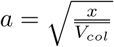

- Phenotypes for scalability and covariance comparison: for the simulated genotype cohort with related individuals, ten independent phenotype sets composed of infinitesimal genetic and observational noise effects with *P ∈ {*10,20,30,50,100*}*, *h*_2_0.2,0.5,0.8 and parameters described in Supplementary Tab. S2 and S3 were simulated;
- phenotypes for calibration analyses: for the both simulated genotype cohorts, phenotypes composed of infinitesimal genetic and observational noise effects with *P* ∈ {10,20,30,50,100}, *h*_2_ ∈ {0.2,0.5,0.8} and parameters described in Supplementary Tab. S2 and S3 were simulated (one phenotype set for each parameter set-up) analyses;
- phenotypes for power analyses: for the simulated genotype cohort with related individuals, phenotypes composed of genetic variant effects, infinitesimal genetic effects, non-genetic covariates and observational noise effects with *P* ∈ {10,50,100}, *h*_2_ ∈ {0.2,0.5,0.8} and parameters described in Supplementary Tab. S2 and S3 were simulated. For each design, 50 random phenotype sets were generated, each with 10 normally distributed non-genetic covariate effects. For each phenotype set, 20 SNP genetic effects were added to a subset of traits (subset sizes in proportion of total number of traits: *a* ∈ {0.2,0.4,0.6,0.8,1}). For all simulations, the mean genetic variance across all traits was kept constant and was fixed at 1% of total phenotypic variance. In total, 3 *h*_2_ × 3 trait sizes ×100 permutations ×5 subset sizes = 4,500 phenotypes were simulated.

### Models

#### Genetic relationship

Population structure and relatedness between individuals can be captured in a genetic relatedness matrix, which accounts for the pairwise genetic similarity between individuals. The relatedness matrices were estimated via ‘plink ‘–make-rel square gz’ option based on an LD-pruned marker set (pruned for variants with *r*^2^ > 0.2) with a window size of 50kb.

#### Multivariate linear models

In the simple multivariate linear model (mvLM), a phenotype 
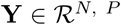
 with *P* traits and *N* samples is modelled as the sum of a genetic fixed effect 
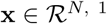
, covariate fixed effects of *K* covariates 
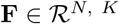
 and i.i.d. residual noise **Ψ**.

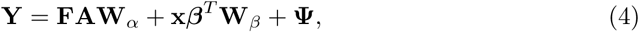

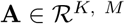
 and 
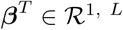
 are the effect size matrix of the covariates and the effect size vector of the genetic variant effects. The trait design matrices **W***_α_* and **W***_β_* allow different scenarios of the cross-trait architecture of the fixed effects on the phenotype. For all analyses in this study, an ‘any effect’ model allowing for independent effects across all traits was chosen, corresponding to **W** = **I**_P_.

#### Multivariate linear mixed models

The multivariate linear mixed model (mvLMM) is an extension of the mvLM through the addition of a genetic random effect **G**:

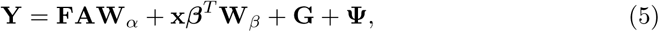

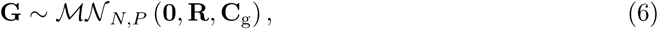

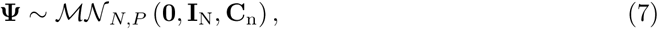

where the random effects **G** and **Ψ** are described by matrix-normal distribution with column covariance **C**_g_ and **C**_n_ and row covariance **R** and **I**_N_, respectively. **C**_g_ and **C**_n_ are the *P* × *P* trait covariance estimates, **R** the *N* × *N* genetic relationship matrix and **I**_N_ a *N* × *N* identity matrix.

#### Univariate linear mixed models

In the univariate linear mixed model (uvLMM), the genetic background and residual noise are modelled as random effects with a scalar estimate for the genetic **G** and noise trait-variance **Ψ**, 
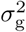
 and 
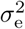
.

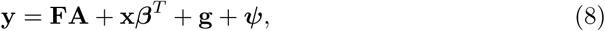

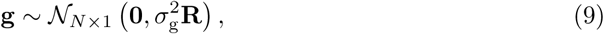

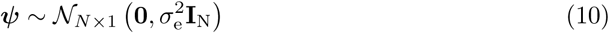

### Covariance comparison

The covariance matrices **C**_g_ and **C**_n_ were estimated via REML and LiMMBo and the goodness of the estimation evaluated based on the root mean square deviation (RMSD) from the true simulated covariance matrices: 
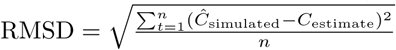
.

### Calibration

LiMMBo and the standard REML approach were used to estimate the trait covariance matrices **C**_g_ and **C**_n_ of the simulated phenotypes without genetic variant effects (null model). **C**_g_ and **C**_n_ were then used as input for a multi-trait GWAS against all genome-wide SNPs. The calibration of the multi-trait GWAS with LiMMBo and standard REML derived trait covariance estimates was visually assessed by comparison against a theoretically expected p-values distribution (quantile-quantile plot). For the comparing the calibration of p-values from the mvLM and mvLMM under the null model, the family-wise error rates (FWERs) of the multi-trait GWAS were estimated by counting the number of tests that exceeded a given threshold divided by the overall number of tests conducted, hence the number of genome-wide SNPs.

### Power analyses

For each of the 20 causal SNPs, a mvLMM and uvLMM against the phenotypes was conducted and the percentage of causal SNPs that were detected i.e. were significantly associated were recorded. The significance was assessed by comparing the p-values obtained from the mvLMM and uvLMM to p-values obtained from mvLMM and uvLMM on 1,000 permutation of the genotypes. For the uvLMM, p-values were adjusted for multiple testing by the number of traits that were tested and the minimum adjusted p-value across all traits for a given SNP recorded. For each SNP, the number of times the (adjusted) p-value of the permutation was less or equal to the observed p-value was recorded and divided by the total number of permutations, yielding an empirical p-value per SNP.

### GWAS of yeast growth traits

#### Data

A publicly available dataset of a yeast cross grown in 46 different conditions [2] was used as a case study to show the feasibility of LiMMBo. It consists of phenotype and genotype data for 1,008 prototrophic haploid *Saccharomyces cerevisiae* segregants derived from a cross between a laboratory strain and a wine strain strain. It contains 11,623 unique genotypic markers for all 1,008 segregants (no missing genotypes) and 46 phenotypic traits. For the phenotyping, segregants were grown on agar plates under 46 different conditions, including different temperatures, pH and nutrient addition (see labels in Supplementary Fig. S8). The traits were defined as end-point colony size in a given condition normalised relative to growth on control medium. Out of the 1,008 segregants, 303 were phenotyped for all 46 traits. Missing values were imputed in segregants that were phenotyped for at least 80% of the traits. The final dataset contained 981 segregants with phenotypes for 41 traits each (a detailed description for the imputation strategy can be found in Supplementary Section 1).

#### Genetic relationship matrix and LD pruning

The genetic relationship matrix of the yeast segregants and different sets of genome-wide SNPs with markers in approximate linkage equilibrium were estimated via plink [43]. Pruning SNPs that are in linkage disequilibrium (LD) was done via ‘–indep-pairwise kb-window 5 0.2’, where the kb-window was varied from 3kb to 100kb. The relationship matrix was estimated via the ‘–make-rel square gz’ option based on an LD-pruned marker (pruned for variants with *r*^2^ > 0.2) set with a window size of 3kb.

#### GWAS

Association of the 11,623 unique genotypic markers with the 41 growth traits was analysed both via mvLMM (Eq. 5), mapping all traits jointly and uvLMM (Eq. 10), mapping each trait individually. In the latter analysis, the p-values obtained were adjusted for multiple testing by the effective number of tests *M_eff_* [44]: 
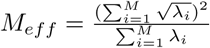
, where *λ* are the eigenvalues of the trait-by-trait correlation matrix. For both analyses, an empirical false discovery rate (FDR) was estimated via permutations, following approaches of previous association studies in yeast crosses [45–47]. With a conservative, theoretical significance threshold of *p*_t_ = 10*^−5^*, at most one SNP is expected to be false positive in a total of *s* = 11,623 SNPs. To find the empirical FDR corresponding to this threshold, *k* = 50 permutations of the genotypes were generated and the LMMs fitted against these permutation. These p-values were used as the empirical p-value distribution and for *p*_t_ = 10*^−5^*, empirical FDRs estimated as FDR_mtGWAS_ = 1.2 × 10^−5^ and FDR_stGWAS_ = 8.6 × 10*^−6^*.

#### Effect size analyses

The following effect size analysis was conducted independently for the effect sizes from the single-trait and multi-trait analyses. All SNPs passing the respective FDR threshold (single-trait or multi-trait) were pruned for LD (*r*^2^ > 0.2, 3kb window; as described above) and location within a gene body (yeast genome assembly: ScerevisaeR64-1-1). The effect size estimates of these SNPs were clustered both across traits and SNPs based on their correlation coefficients via *pvclust* [27] (number of iterations: 50,000 for traits and 10,000 for SNPs). *pvclust* yields bootstrap-based p-values as a measure for the stability of a given cluster. Clusters with *p* < 0.05 were considered significant and extracted via *pvpick*, setting max.edge to FALSE.

## Acknowledgments

We thank Danilo Horta for his help with the module development.

## Supporting information

### 1 Supplementary Data

#### LMM inference schemes

Supplementary Tab. S1 summarises commonly used frameworks and describes their computational complexity^1^.

**Table S1.**
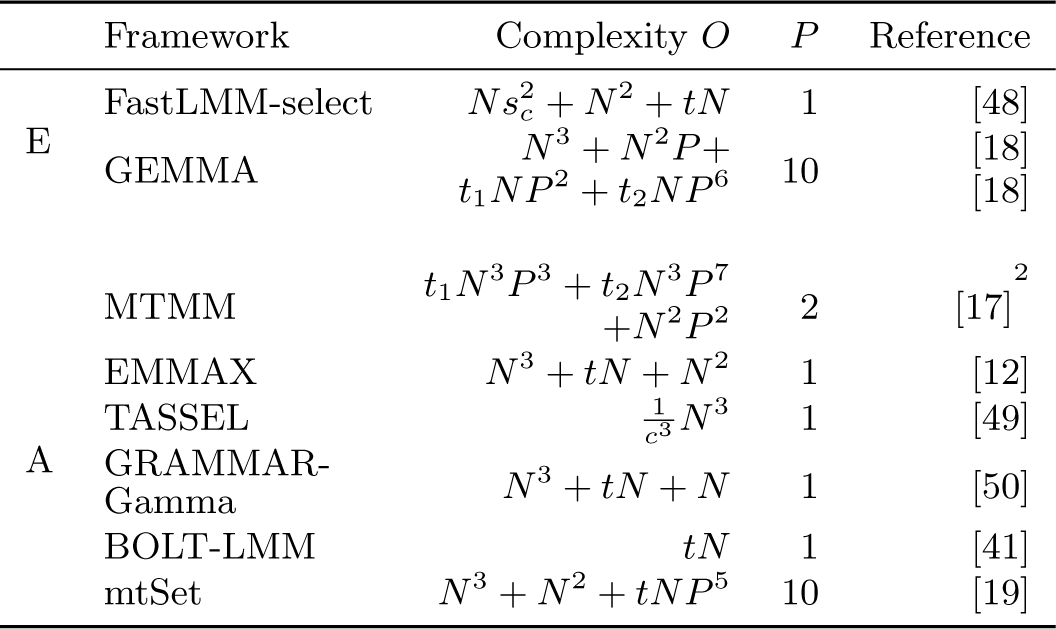
Linear mixed model frameworks for genetic association studies. A list of popular LMM frameworks, grouped by their usage of covariance estimates when fitting the alternative model (first column: E: exact, A: approximate). The complexity describes the complexity for fitting a single LMM as specified in the original publication or summarised elsewhere, as indicated by the footnotes. *P* indicates the trait size that the model was designed for (according to the original publication). Models with specific parameters are described in more detail in the text (FaST-LMM-select and TASSEL). *N*: number of samples; *s_c_*: number of SNPs used for singular value decomposition; *c*: compression factor with 
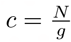
 for *g* individuals per group; *t*, *t*_1_and*t*_2_: average number of iterations needed to find parameter estimates. GRAMMAR-Gamma, FaST-LMM-select: *t* steps of the Brent’s algorithm; GEMMA, MTMM: *t*_1_ steps of the EM algorithm, *t*_2_ steps of the NR algorithm; BOLT-LMM: *t* steps of the variational Bayes and conjugate gradients; TASSEL: *t* steps of the ProcMixed algorithm in SAS; mtSet: *t* steps of the L-FBGS.

Among the exact methods, FaST-LMM-select reduces the complexity best in terms of sample size by selecting the number of SNPs to use for the estimation of the relatedness matrix. However, it can only be applied in univariate analyses while MTMM and GEMMA extend to multivariate cases. BOLT-LMM scales best with increasing samples sizes in the group of approximate tests, by directly using the genotypes and not computing or storing the relatedness matrix. All other methods have an upfront *O*(*N*^3^) operation for the eigendecomposition of the relatedness matrix. TASSEL reduces this complexity based on grouping of the samples and thereby effectively reducing the size of the relatedness matrix

#### Simulations

##### Genotypes

The synthetic genotypes were generated based on real genotype data from four European ancestry populations of the 1000 Genomes (1KG) Project (populations: CEU, FIN, GBR, TSI) [24] (see main methods). The genetic relatedness matrices and scatter plots of the first two principal components for each cohort are shown in Supplementary. S1.

**Fig S1.**
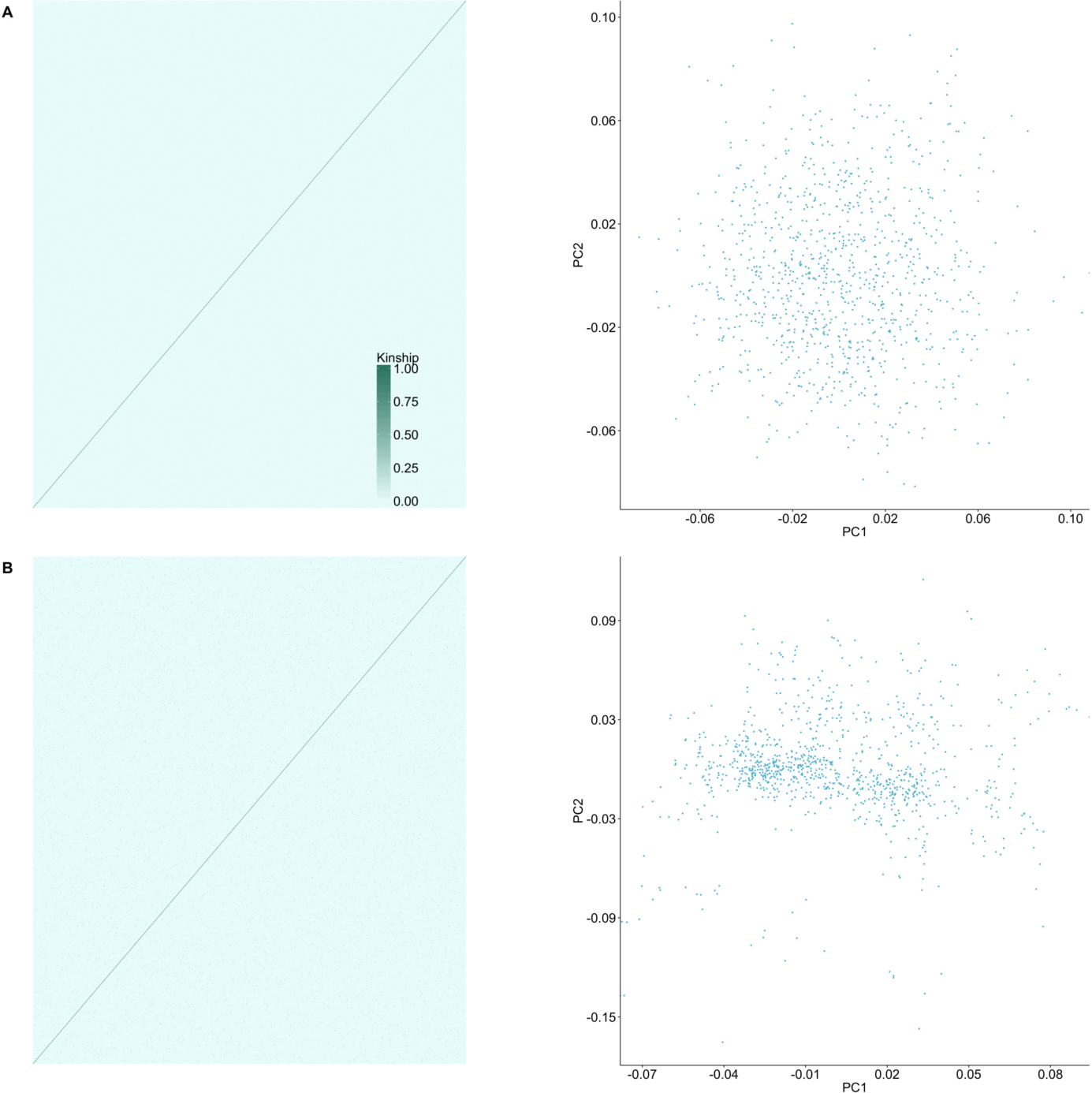
Kinship matrices and principal components of two simulated European ancestry cohorts. The genotypes were simulated based in genotype data from four European ancestry population. Depending on the choice and number of ancestors *a* for the sampling of chromosomes to simulate an individuals genotype, cohorts with differing levels of population and relatedness structure will arise. A. unrelated individuals *a* = 10. B. related individuals *a* = 2.

#### Phenotypes

**Table S2.** Parameters for phenotype simulation. For each, the total variance is the sum of both effect variances and has to add to 1. Each component has a certain percentage of its variance that is shared across traits, while the rest is independent.

|  |  | variance explained | shared | independent |
| --- | --- | --- | --- | --- |
| genetic effects | total | $h_2$ | | |
| | SNP | $h_2 h_2^s$ | $\theta$ | $1-\theta$ |
| | infinitesimal | $h_2 h_2^g$ | $\eta$ | $1-\eta$ |
| noise effects | total | $(1-h_2)$ | | |
| | covariates | $(1-h_2)\delta$ | $\gamma$ | $1-\gamma$ |
| | observational noise | $(1-h_2)(1-\delta)$ | $\alpha$ | $1-\alpha$ |

**Table S3.** Parameter values of simulated phenotypes for assessing scalability, calibration and power. *P* are the different traitset sizes that were simulated. The parameters that follow are described in Tab. S2 and specify the variance explained by each of the phenotype components. A variance explained equals zero means that this component was not simulated and corresponding non-applicable variance terms are designated with ‘-’.

| Parameter | Parameter values |  |  |
| --- | --- | --- | --- |
|  | Scalability | Covariance comparison | Calibration |
| Genotypes | relatedNoPopstructure | relatedNoPopstructure | relatedNoPopstructure<br>unrelatedPopstructure |
| $P$ | 10, 20, ..., 100 | 10, 20, 30, 50, 100 | 10, 20, 30, 50, 100 |
| $h_2^s$ | 0 | 0 | 0 |
| $h_2^g$ | 1 | 1 | 1 |
| $h_2$ | 0.8, 0.5, 0.2 | 0.8, 0.5, 0.2 | 0.8, 0.5, 0.2 |
| $(1-h_2)\delta$ | 0 | 0 | 0 |
| $(1-h_2)(1-\delta)$ | 1 | 1 | 1 |
| $(1-h_2)$ | 0.2, 0.5, 0.8 | 0.2, 0.5, 0.8 | 0.2, 0.5, 0.8 |
| $\theta$ | - | - | - |
| $\eta$ | 0.8 | 0.8 | 0.8 |
| $\gamma$ | - | - | - |
| $\alpha$ | 0.8 | 0.8 | 0.8 |

| Parameter | Power |
| --- | --- |
| Genotypes | relatedNoPopstructure |
| $P$ | 10, 50, 100 |
| $h_2^s$ | 0.05, 0.02, 0.0125 |
| $h_2^g$ | 0.95, 0.98, 0.9875 |
| $h_2$ | 0.8, 0.5, 0.2 |
| $(1-h_2)\delta$ | 0.4 |
| $(1-h_2)(1-\delta)$ | 0.6 |
| $(1-h_2)$ | 0.2, 0.5, 0.8 |
| $\theta$ | 0.6 |
| $\eta$ | 0.8 |
| $\gamma$ | 0.6 |
| $\alpha$ | 0.8 |

#### Model calibration: linear mixed model versus simple linear model

We compared the calibration of mtGWAS using a mvLMM to the calibration of a simple multivariate linear model (LM, Eq. 4). The LM does not require the variance decomposition into different random effects, i.e. avoids the computational bottleneck, but simply uses principal components of the genotypes as fixed effects to adjust for population structure. Supplementary Tab. S4 shows the type I error estimates for both GWAS approaches. The LMM (Eq. 5) approach performs well across all trait sizes and thresholds of significance. In contrast, the LM is poorly calibrated and clearly demonstrates the difficulty of adjusting for population structure via fixed effects in highly structured populations.

**Table S4.** Type I error estimates for mtGWAS. Type I errors estimates for mvLMM and mvLM across all genome-wide SNPs for three trait set sizes assessed at two different levels of significance. For the mvLMM, covariance estimates were derived via LiMMBo. In the mvLM, population structure was adjusted for via the first ten PCs of the genotype data. The mvLMM controls well for Type I errors at both thresholds, while the mvLM leads to inflated test-statistics.

| Traits | Significance level | Type I Error estimates |  |
| --- | --- | --- | --- |
|  |  | mvLM | mvLMM |
| 10 | 5.00E-05 | 2.17E-03 | 4.51E-05 |
|  | 5.00E-08 | 2.32E-05 | <5E-08 |
| 50 | 5.00E-05 | 1.09E-02 | 1.93E-05 |
|  | 5.00E-08 | 2.08E-04 | <5E-08 |
| 100 | 5.00E-05 | 3.17E-02 | 2.28E-05 |
|  | 5.00E-08 | 1.01E-03 | 4.56E-08 |

#### Power analyses

**Fig S2.**
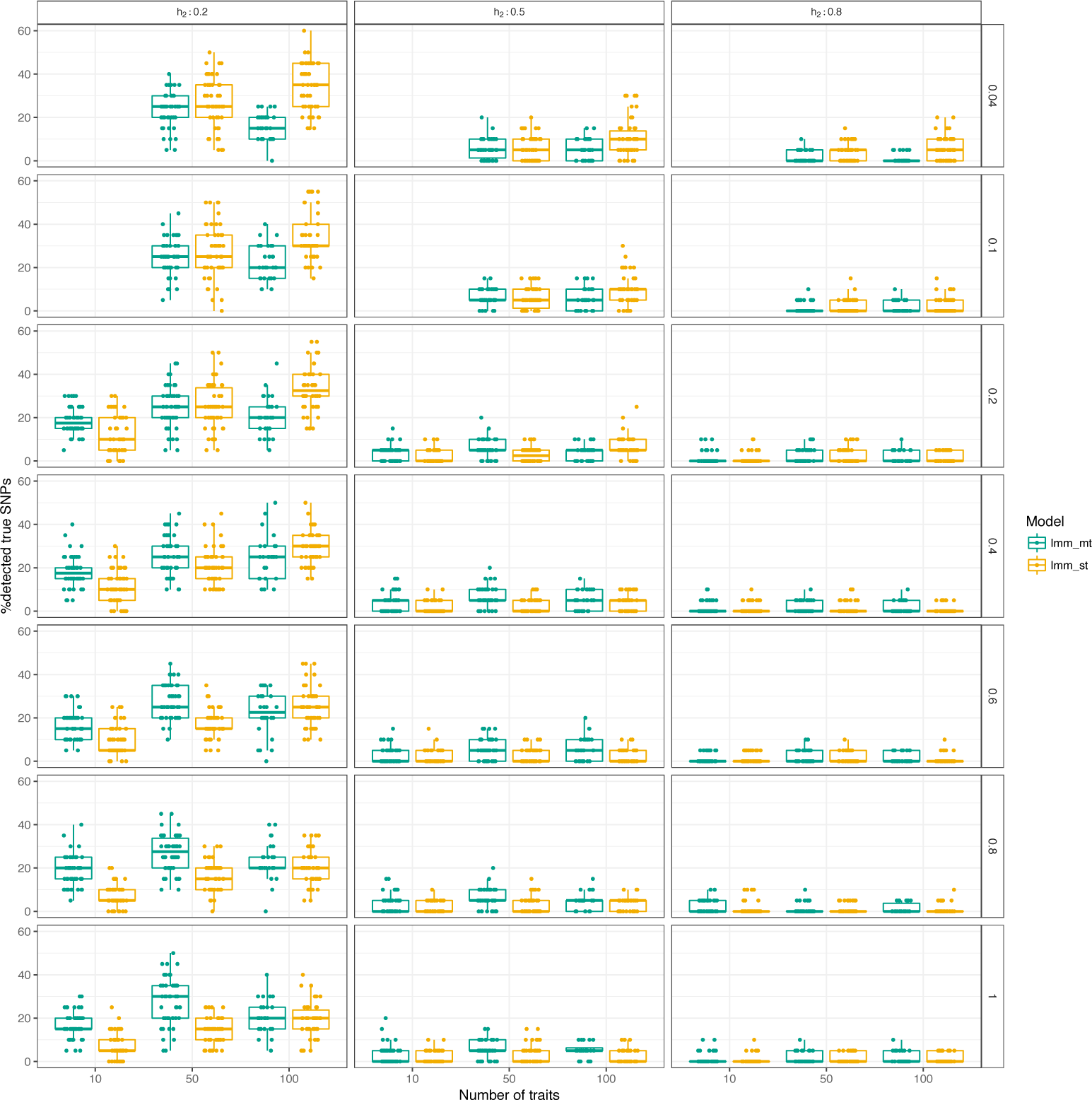
All parameter combinations of power comparison for multivariate and univariate LMM of high-dimensional phenotypes. Each panel shows the influence of one simulation parameter on the power to detect the causal SNPs: rows show the influence of the proportion of traits affected by the genetic variant effects, columns show the influence of the overall phenotypic variance explained by genetics.

#### Multi-trait association study of yeast growth traits

**Table S5.** Comparison of significant loci in single-trait and multi-trait GWAS. In column ‘All SNPs’, the absolute number of SNPs beyond the significance threshold for multi-trait and single-trait GWAS as well as their ratio (multitrait:singletrait) are depicted. In order to limit the potential bias in the counting of the loci (different degrees of linkage disequilibrium (LD) for different loci and genotyping parameters), the genome-wide SNPs were LD pruned and the ratio of significant SNPs determined for five different LD window sizes. The maximal LD window covering between 6% (chromosome 4) and 43% (chromosome 1) of total chromosome length.

| | All SNPs | LD pruned with $r^2 \geq 0.2$ | | | | |
| --- | --- | --- | --- | --- | --- | --- |
|  |  | 3kb | 10kb | 30kb | 50kb | 100kb |
| NrSNPs | 11623 | 4105 | 1028 | 264 | 161 | 107 |
| multitrait | 1132 | 384 | 101 | 24 | 15 | 9 |
| singletrait | 695 | 275 | 72 | 20 | 13 | 7 |
| multitrait:singletrait | 1.63 | 1.4 | 1.4 | 1.2 | 1.15 | 1.29 |

**Fig S3.**
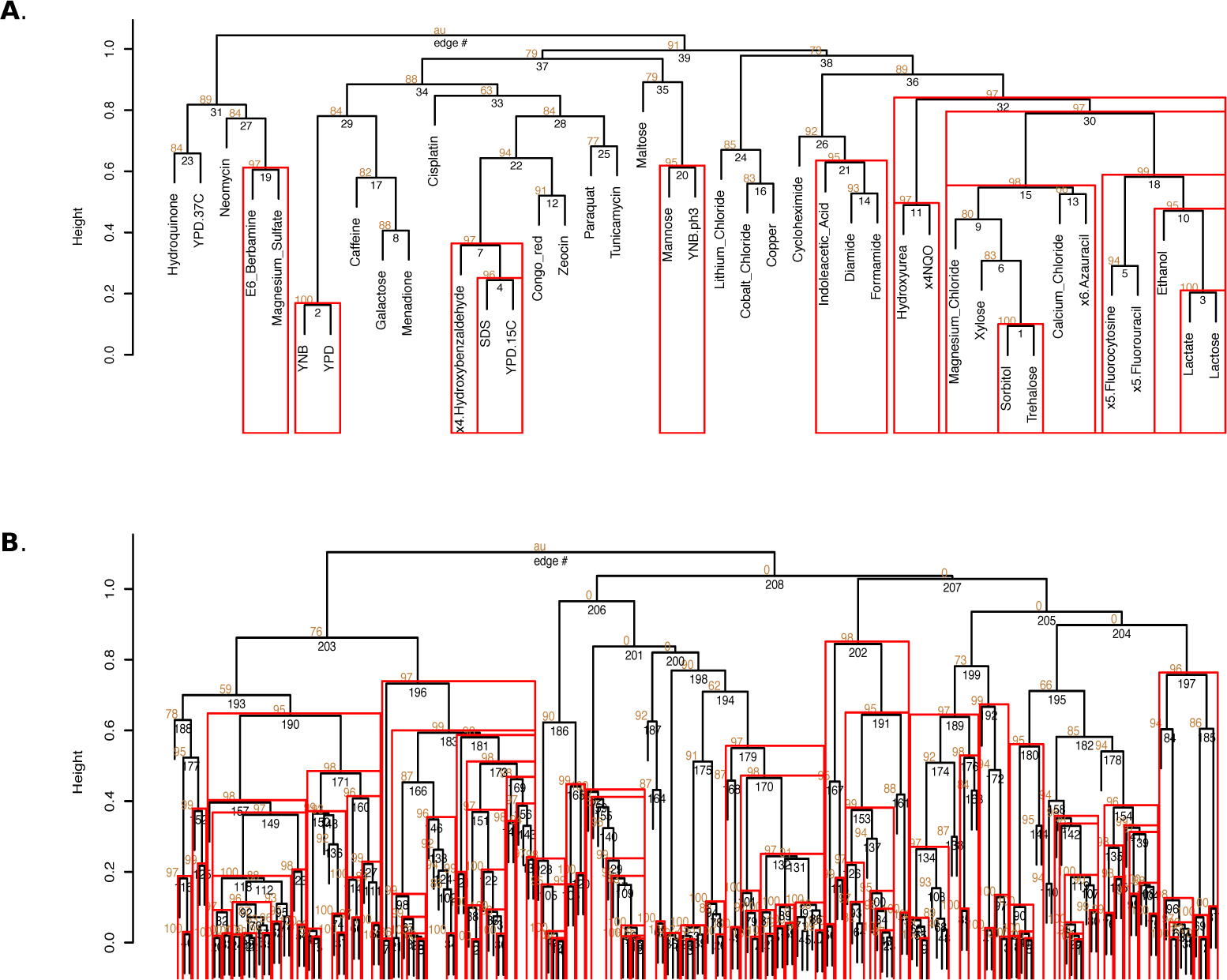
Dendrograms of hierarchical cluster analyses of multi-trait effect size estimates. The effect size estimates of significant SNPs from the multi-trait GWAS were filtered for SNPs located in a gene body and subsequently LD-pruned (3kb window, *r*^2^ > 0.2). The resulting set of 210 SNPs was independently clustered across traits and SNPs, using a hierarchical clustering based on the correlation of the effect size estimates. The clustering was repeated 50000 (for traits) and 10000 (for SNPs) via pvclust [27], allowing for the estimation of stable clusters – recurring clusters, approximately unbiased (‘au’) values of greater or equal to 0.95. ‘au’ values are depicted on the vertices of the dendrogram. Stable clusters with vertices of *au* ≥ 0.95 are enclosed by a red box (generated with *pvclust::pvpickPlot* and *pvclust::pvpickRect*). Some stable clusters enclose other, smaller stable clusters (for instance right-most large cluster in A) A. Clustering of growth traits based on effect size estimates: 14 stable clusters B. Clustering of SNPs based on effect size estimates (SNP labels are omitted): 136 stable clusters.

**Fig S4.**
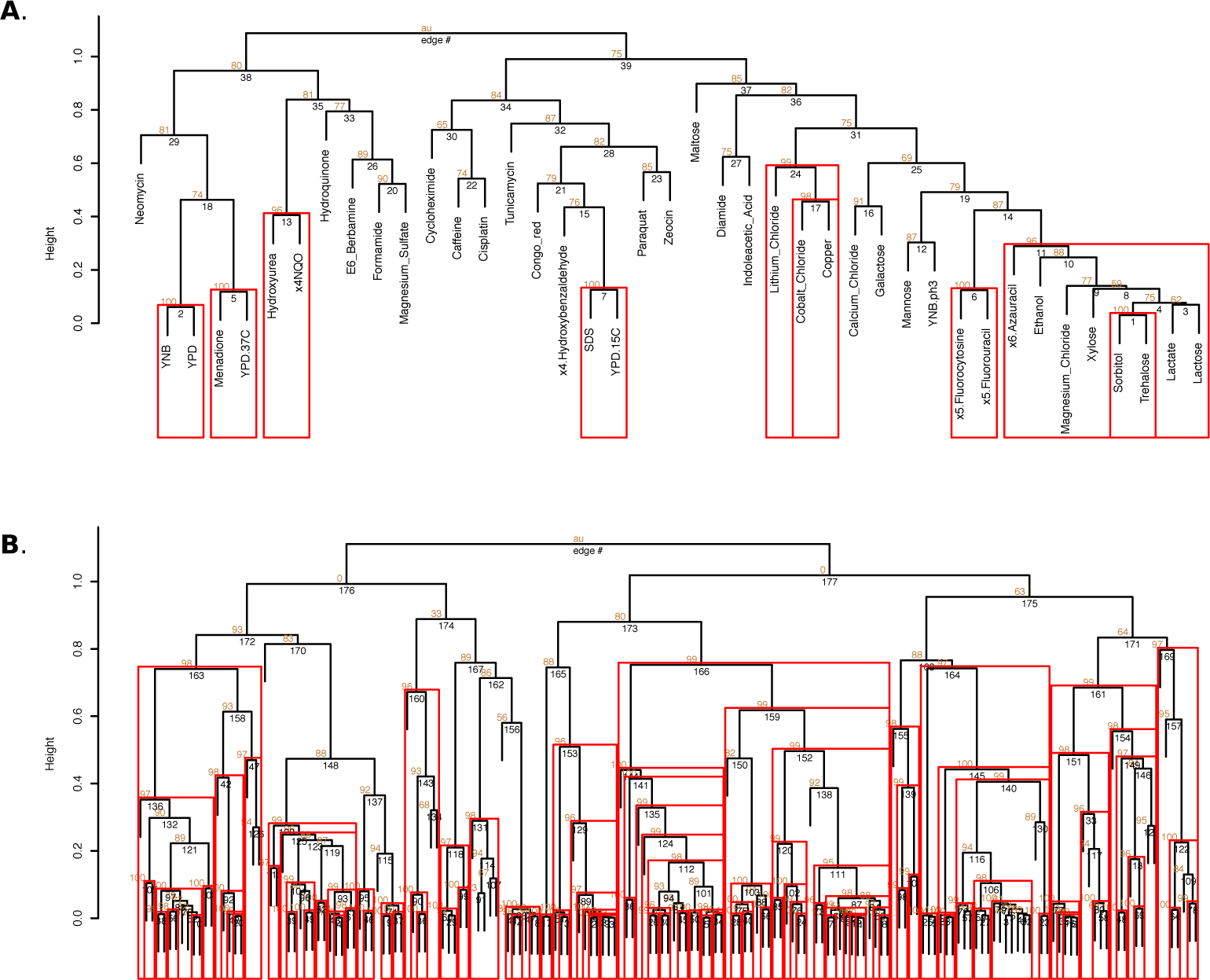
Dendrograms of hierarchical cluster analyses of single-trait effect size estimates. The effect size estimates of significant SNPs from the single-trait GWAS were filtered for SNPs located in a gene body and subsequently LD-pruned (3kb window, *r*^2^ > 0.2). The resulting set of 179 SNPs was independently clustered across traits and SNPs, using a hierarchical clustering based on the correlation of the effect size estimates. The clustering was repeated 50000 (for traits) and 10000 (for SNPs) via pvclust [27], allowing for the estimation of stable clusters – recurring clusters, approximately unbiased (‘au’) values of greater or equal to 0.95. ‘au’ values are depicted on the vertices of the dendrogram. Stable clusters with vertices of *au* ≥ 0.95 are enclosed by a red box (generated with *pvclust::pvpickPlot* and *pvclust::pvpickRect*). Some stable clusters enclose other, smaller stable clusters (for instance right-most large cluster in A) A. Clustering of growth traits based on effect size estimates: 9 stable clusters B. Clustering of SNPs based on effect size estimates (SNP labels are omitted): 117 stable clusters.

**Fig S5.**
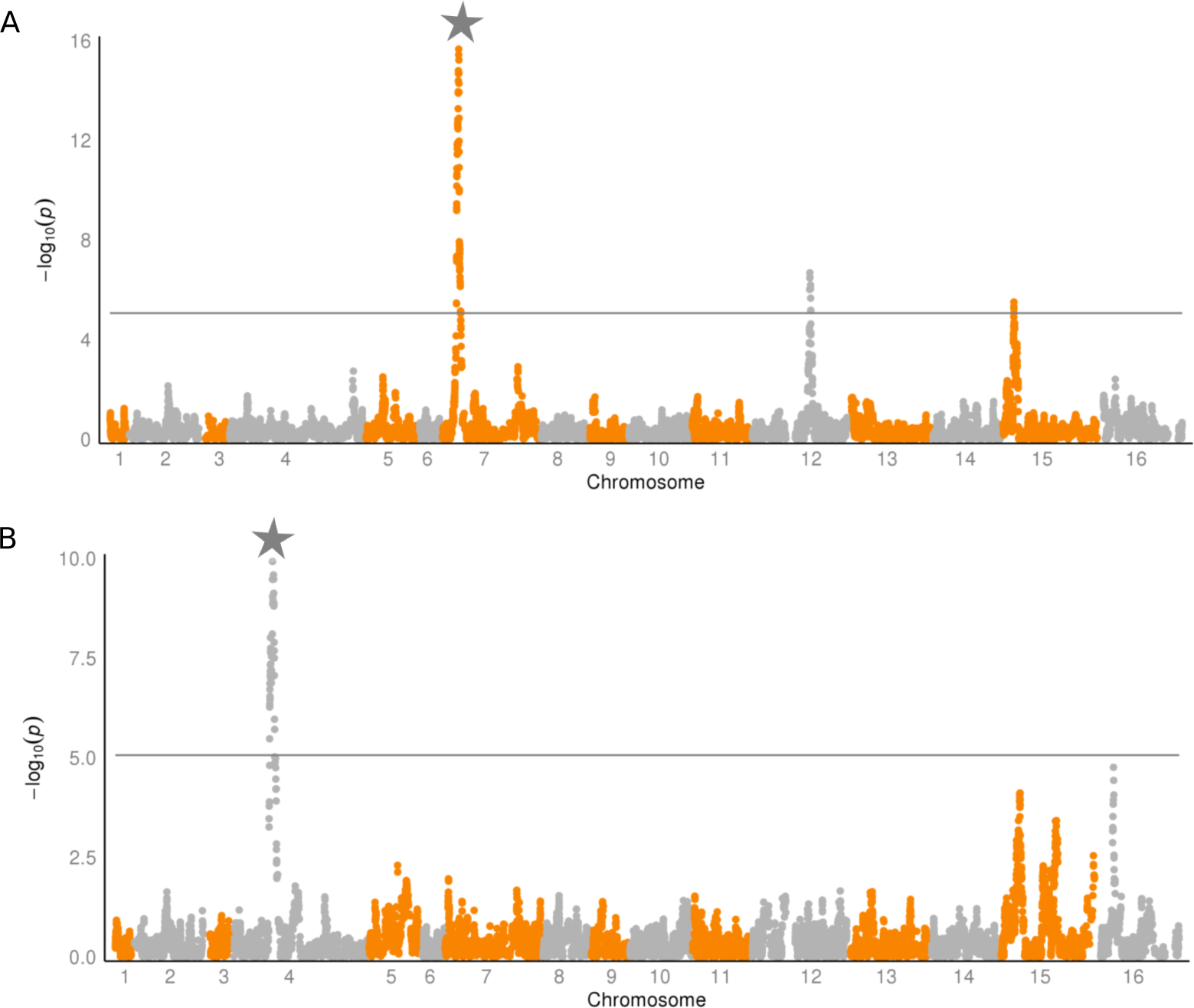
Manhattan plot of traits with strong single-trait associations. Single-trait GWAS of A. magnesium sulfate and B. hydroquinone. The loci marked with a grey star are only found for these two traits and cannot be detected in the multi-trait GWAS (Fig. 5), pointing to purely single-trait association that is burdened by the multi-trait testing based on 41 degrees of freedom. The p-values were adjusted for multiple testing by the effective number of tests (*M*_eff_ = 33). The significance line is drawn at the empirical FDR_stGWAS_ = 8.6 *×* 10*^−6^*.

**Fig S6.**
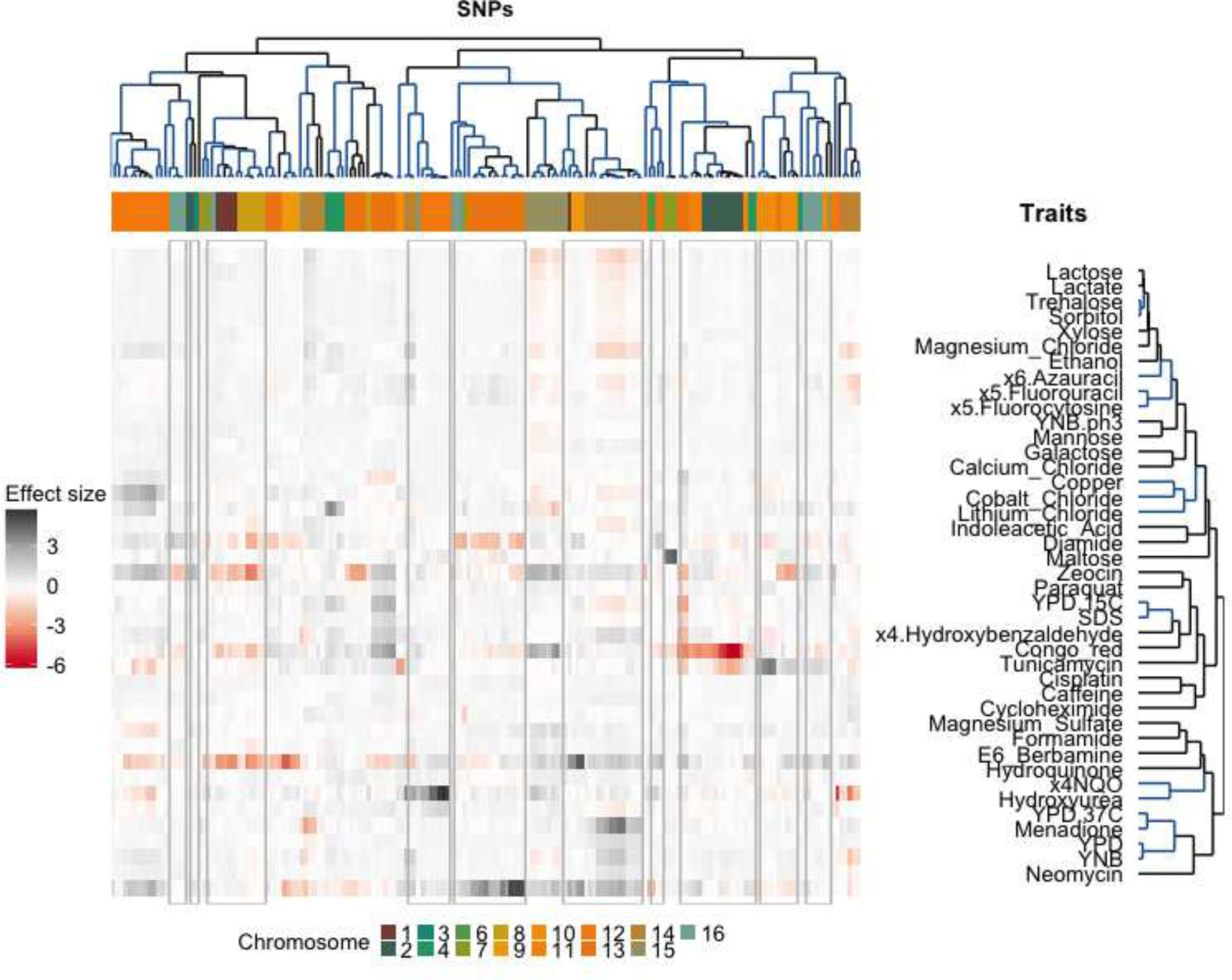
Single-trait GWAS effects size estimates. Significant SNPs (695 SNPs; adjusted for multiple testing, passing the threshold FDR_stGWAS_ = 8.6 × 10^−6^) of the single-trait GWAS for the 41 yeast growth traits were LD-pruned (3kb window, *r*^2^ > 0.2) and filtered for SNPs located within a gene body (final SNP count: 179). The effect size estimates of these SNPs were clustered by loci and traits (both hierarchical, average-linkage clustering of Pearson correlation coefficients). Traits and SNPs in stable clusters (pvclust*p* < 0.05) are marked in blue.

#### Imputation of missing phenotypes in yeast dataset

The dataset consists of phenotype and genotype data of 1,008 prototrophic haploid *Saccharomyces cerevisiae* segregants derived from a cross between a laboratory strain and a wine strain strains [2]. It contains 11,623 unique genotypic markers obtained via short-read sequencing for all 1,008 segregants (no missing genotypes). For phenotyping, segregants were grown on agar plates under 46 different conditions, including different temperatures, pH and nutrient addition (see labels in Supplementary Fig. S8). The phenotypes were defined as end-point colony size normalised relative to growth on control medium. In the following, a trait is defined as the normalised growth size in one condition.

Out of the 1,008 segregants, 303 were phenotyped for all 46 traits. Missing phenotypes are not evenly distributed with some traits such as cobalt chloride being present for almost all samples while others such as sorbitol or raffinose are lacking in more than a third of the samples. The distribution of trait missingness across all samples is depicted in Supplementary Fig. S7A.

The LMM framework relies on all samples being fully genotyped and phenotyped and does not accept missing values. In order to use the largest possible subset of the data, imputation strategies for the missing phenotypes were sought. We used the subset of 303 fully phenotyped samples to determine traits suitable for imputation. We simulated data with a similar pattern of missingness as observed in the original dataset by subsampling the full dataset to the subset size and overlaying the observed missingness pattern onto the subset of 303 fully phenotyped samples. The resulting pattern is depicted in Supplementary Fig. S7B. Similar results in frequencies of fully phenotyped samples and combination of missing/non-missing traits can be observed when comparing it to the original frequencies and patterns (Supplementary Fig. S7A). We chose the MICE framework [52] with PMM as the imputation method to determine the most suitable imputation parameter settings in the simulated dataset which would then be applied to impute the real missing values in the full dataset. The predictor variables for each trait were determined based on their pair-wise Spearman’s rank correlation coefficient *ρ* with all other traits in the dataset (Supplementary Fig. S8). In addition, only predictor traits that had been measured in at least 20% of the samples in the dataset were considered. Different predictor variable set-ups were examined based on increasing thresholds for Pearson’s correlation coefficient: *r*^2^0,0.1,0.2,0.3.

Further parameters for MICE are the number of multiple iterations *m* (set to *m* = 20) and the number of iterations *maxit* (set to *maxit* = 30). For each predictor set-up, MICE was initiated with the same seed for the random number generator to ensure comparability. The goodness of the imputation was evaluated by computing the correlation of the imputed values (averaged across iterations *m*) to the experimentally observed ones. Traits where the imputed values correlated to the original ones by more then 95% in at least one of the predictor set-ups were retained in the analysis. For five traits (cadmium chloride, hydrogen peroxide, raffinose, YNB:ph8, YPD:4C), no suitable predictors could be determined and these were excluded from further analyses (Supplementary Fig. S9, red labels). For each trait, the predictor scheme that yielded the highest correlation between the imputed and observed data was chosen for the imputation of missing values in the full dataset. Missing values were imputed in segregants that were phenotyped for at least 80% of the traits. The final dataset contained 981 segregants with phenotypes for 41 traits each.

**Fig S7.**
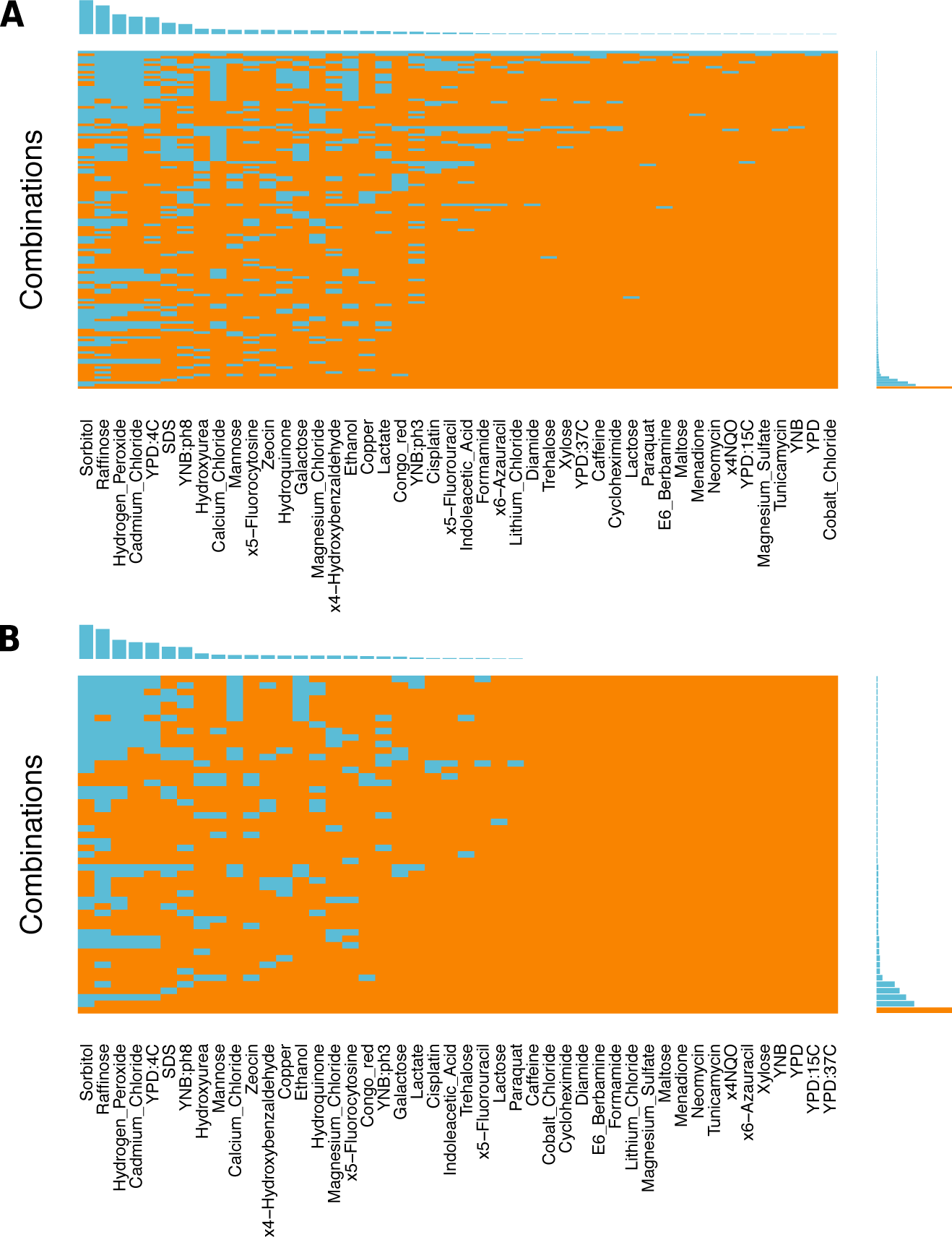
Frequencies and distributions of missing values in the yeast phenotype data. In both panels, the aggregation plot (middle) depicts all existing combinations of missing (blue) and non-missing (orange) values in the traits. The bar chart on its right shows the frequencies of occurrence of the different combinations. The histogram on the top shows the frequency of missing values for each trait (R Package: *VIM* [51]). A. The full dataset contains normalised colony sizes for growth in 46 different conditions of 1,008 genotyped yeast segregants. 306 segregants are fully genotyped (bar chart, orange bar). B. Fully-phenotyped dataset of 306 segregants with simulated missing values based on the observed missingness pattern for the entire pool of 1,008 segregants. Generated via R function *VIM::aggr*.

**Fig S8.**
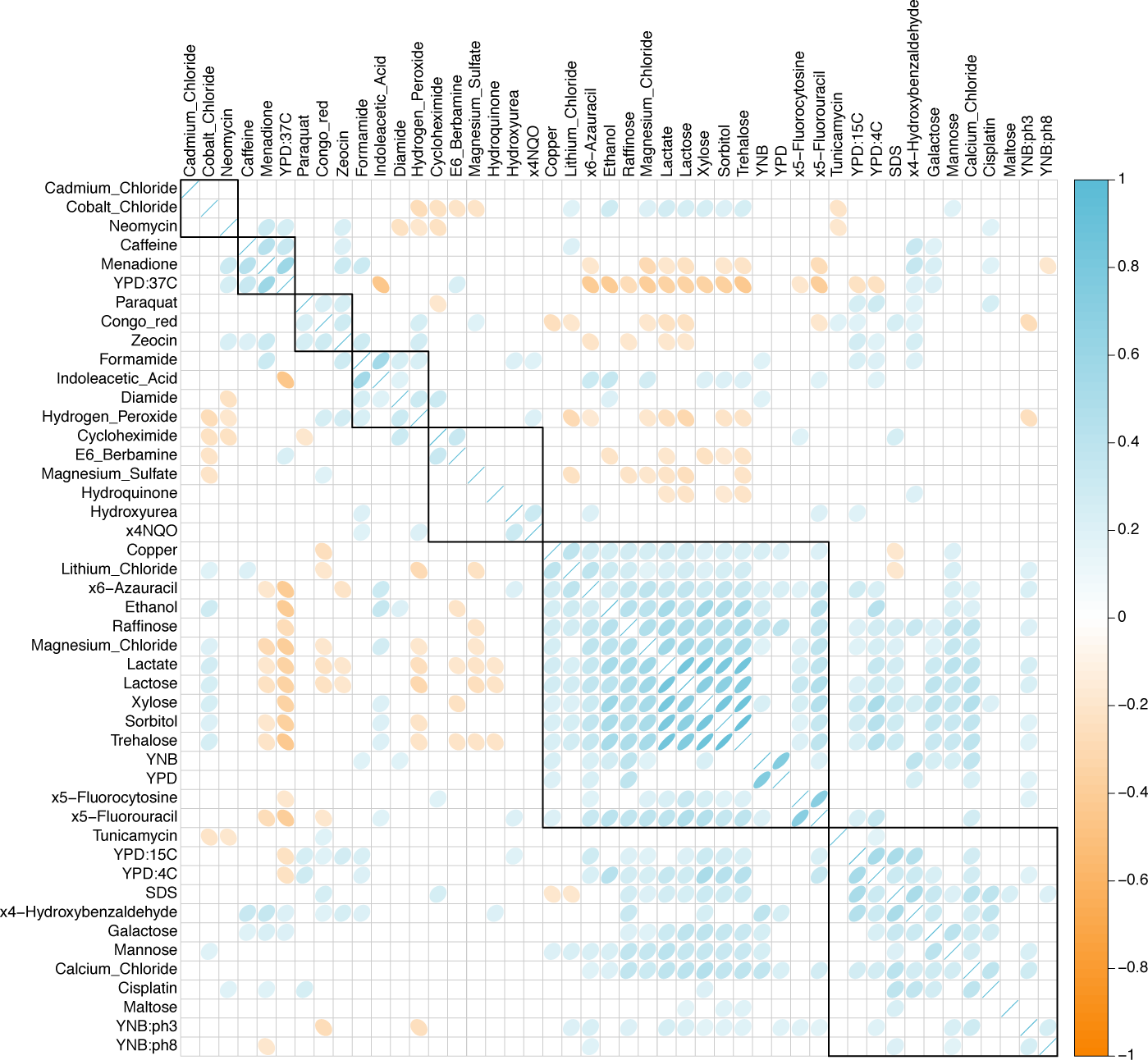
Pair-wise correlations of 46 growth traits in *Saccharomyces cerevisiae*. For each trait pair, Pearson’s correlation coefficient *r*^2^ and the p-values of the correlation were computed. The p-values were adjusted for multiple testing according to Benjamini and Hochberg’s method [53]. The strength and the direction of significant correlations (*p* < 0.05) are depicted above. Non-significant correlations are left blank. The traits are clustered based on complete-linkage clustering of (1*r*^2^) as distance measurement (R Package: *corrplot*).

**Fig S9.**
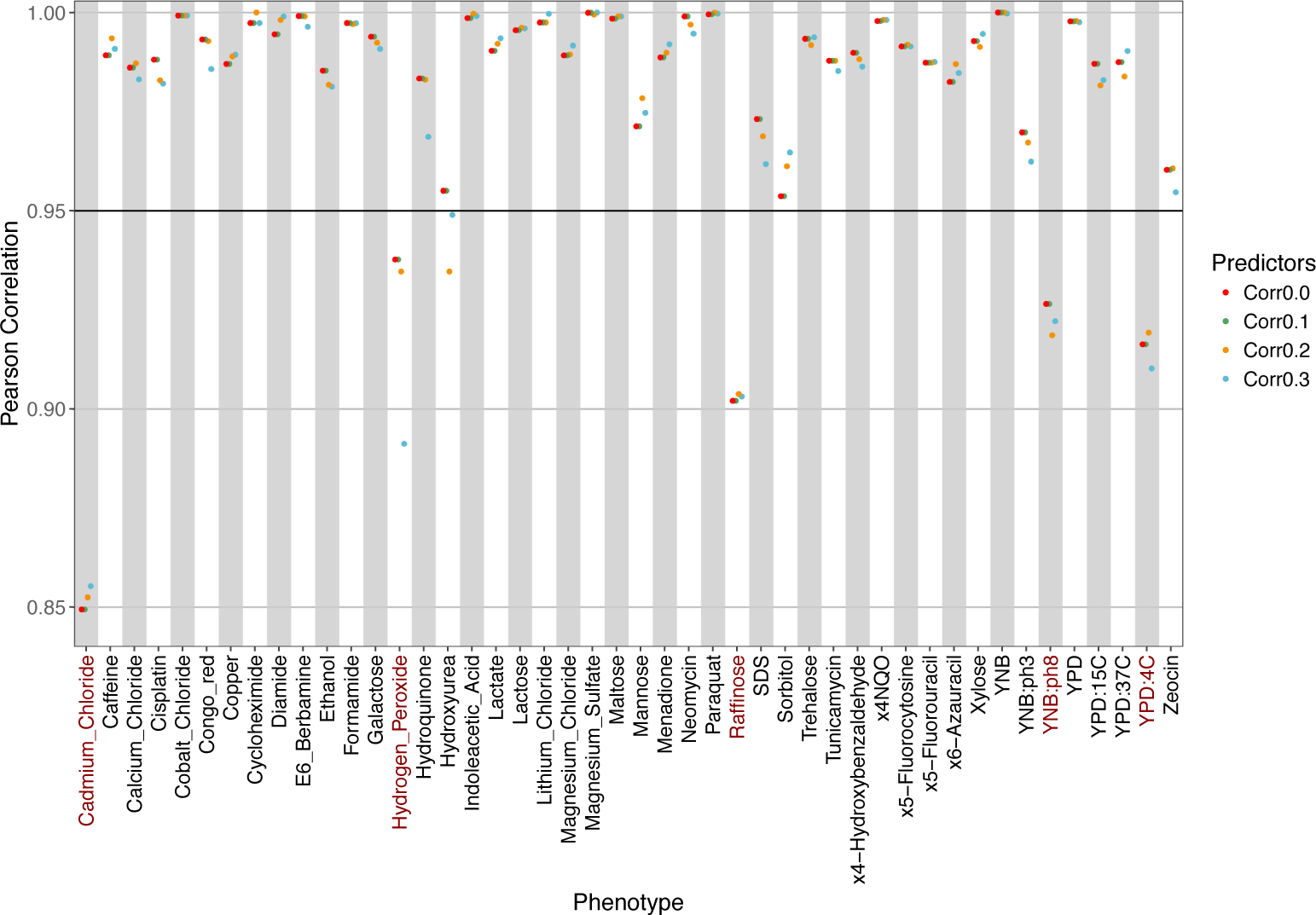
Correlation between imputed and experimentally observed trait values. In the subset of 306 fully phenotyped samples, missing values were introduced and subsequently imputed via MICE. Different predictor sets were tested, differing in the predictors traits included. Sets were constructed based on different Spearman’s rank correlation coefficient: traits were considered predictors if their correlation with the target trait was greater than a given threshold. For each predictor setup *ρ* 0,0.1,0.2,0.3, *m* = 20 multiple imputations and *maxit* = 30 iterations of MICE were conducted. The goodness of the imputation was evaluated by computing the correlation of the imputed values (averaged across iterations *m*) to the experimentally observed ones. Traits with at least one correlation greater than the 0.95 (black vertical line) were retained in the dataset. For traits labelled in red, the imputation was considered to be unreliable and the traits were excluded from further analyses (R Package: *mice* [52]).

## Footnotes

1 The computational complexity and algorithms for the GCTA implementations [15] of multivariate genetic variance estimation [16] and LMM for association testing [13] could not be found in the original publications and are therefore not listed

